# REPTOR/CREBRF encode key regulators of muscle energy metabolism

**DOI:** 10.1101/2021.12.17.473012

**Authors:** Pedro Saavedra, Phillip A. Dumesic, Yanhui Hu, Patrick Jouandin, Richard Binari, Sarah E. Wilensky, Elizabeth Filine, Jonathan Rodiger, Haiyun Wang, Bruce M. Spiegelman, Norbert Perrimon

**Author notes:** Corresponding authors: Pedro Saavedra, Norbert Perrimon.

## Abstract

Metabolic flexibility of muscle tissue describes the capacity to use glucose or lipids as energy substrates and its disruption is associated with metabolic dysfunction. Cancer-induced cachexia is a metabolic syndrome linked with muscle wasting, changes in muscle energy metabolism and lower life expectancy in cancer patients. The molecular mechanisms driving metabolic changes in muscle, however, are poorly characterized. Here, using a Drosophila model of systemic metabolic dysfunction triggered by yorkie-induced gut tumors, we identify the transcription factor REPTOR as a key regulator of energy metabolism in muscle. We show that *REPTOR* is upregulated in muscles of adult flies with gut yorkie-tumors, where it is necessary to modulate glucose metabolism. *REPTOR* expression in muscles is induced by ImpL2, a tumor-derived insulin binding protein that reduces systemic insulin signaling, or by nutritional restriction. Further, in vitro and in vivo studies indicate that high activity of REPTOR is sufficient to increase glucose content, transcriptionally repress phosphofructokinase and increase mitochondrial respiration. Consistent with the fly studies, higher levels of CREBRF, the mammalian ortholog of REPTOR, reduce glycolysis in mouse myotubes while promoting an oxidative phenotype. Altogether, our results implicate REPTOR/CREBRF as key regulators of muscle metabolism and metabolic flexibility that share a conserved function as repressors of glycolysis and promoters of oxidative phosphorylation.

---

Metabolic flexibility refers to the ability of tissues like muscle to use glucose or lipids as energy substrates depending on their availability(*1-3*). After feeding, an increase in glucose levels reduces fatty acid oxidation in muscle, whereas nutritional restriction increases the supply of fatty acids to the muscle and suppresses glucose usage(*4, 5*). Disruption of this flexibility, however, is frequently observed in the context of overwhelming abundance of either glucose or lipids, and associates with insulin resistance, obesity and disruption of muscle energy metabolism(*2, 3, 6*). It is therefore important to identify genes that regulate energy substrate selection by muscles, and that can have therapeutical potential in the context of metabolic syndromes. One such syndrome is cancer induced-cachexia, a complex systemic metabolic dysfunction observed in cancer patients that severely affects skeletal muscle and adipose tissue*(7-9)*. Skeletal muscle of cancer patients displays impaired glucose metabolism and insulin resistance(*10*), whereas the adipose tissue often exhibits higher lipolysis(*11*). Mitochondrial degeneration and dysfunction during cachexia has also been associated with muscle wasting and loss of sarcomere integrity(*12, 13*), suggesting a link between energy metabolism and muscle wasting. In *Drosophila*, two models of tumor-induced organ wasting have been described(*14, 15*). Both tumor models secrete an insulin binding protein (IBP), ImpL2(*16, 17*), that reduces insulin signaling in peripheral tissues, induces organ wasting and systemic metabolic dysfunction. Here, we use the *yorkie* (*yki*) gut model of tumors*(14)* (*Esg*^*[TS]*^*>yki*^*[S3A]*^) to identify novel genes that regulate muscle metabolism by focusing on the flight muscles of the adult fly thorax.

In cancer patients with cachexia, metabolic changes in muscle include insulin resistance and changes in glucose/fatty acid utilization(*18*), and increased lipolysis prior to muscle wasting(*19*). To study how gut *yki*-tumors affect muscle energy metabolism in adult flies, we measured triglyceride (TAG) and glucose content in *Esg*^*[TS]*^*>yki*^*[S3A]*^ thoraces over time. The adult thorax is mainly composed of muscle fibers that control wing and leg movement but also has portions of fat body, the fly adipose tissue (Fig. 1A). TAG levels slowly decreased with aging in control thoraces but *Esg*^*[TS]*^*>yki*^*[S3A]*^ thoraces displayed a sharp reduction in TAG content after 8 days of tumor induction (Fig. 1B), suggesting increased TAG breakdown(*14*). Conversely, we observed a progressive increase in tissue glucose content, from 8 to 14 days after tumor induction (Fig. 1C), which may indicate a reduction in glucose usage. Taken together, these results suggest that *Esg*^*[TS]*^*>yki*^*[S3A]*^ flies display changes in TAG and glucose utilization similar to what is observed in cancer patients with muscle metabolic changes.

**Fig. 1:**
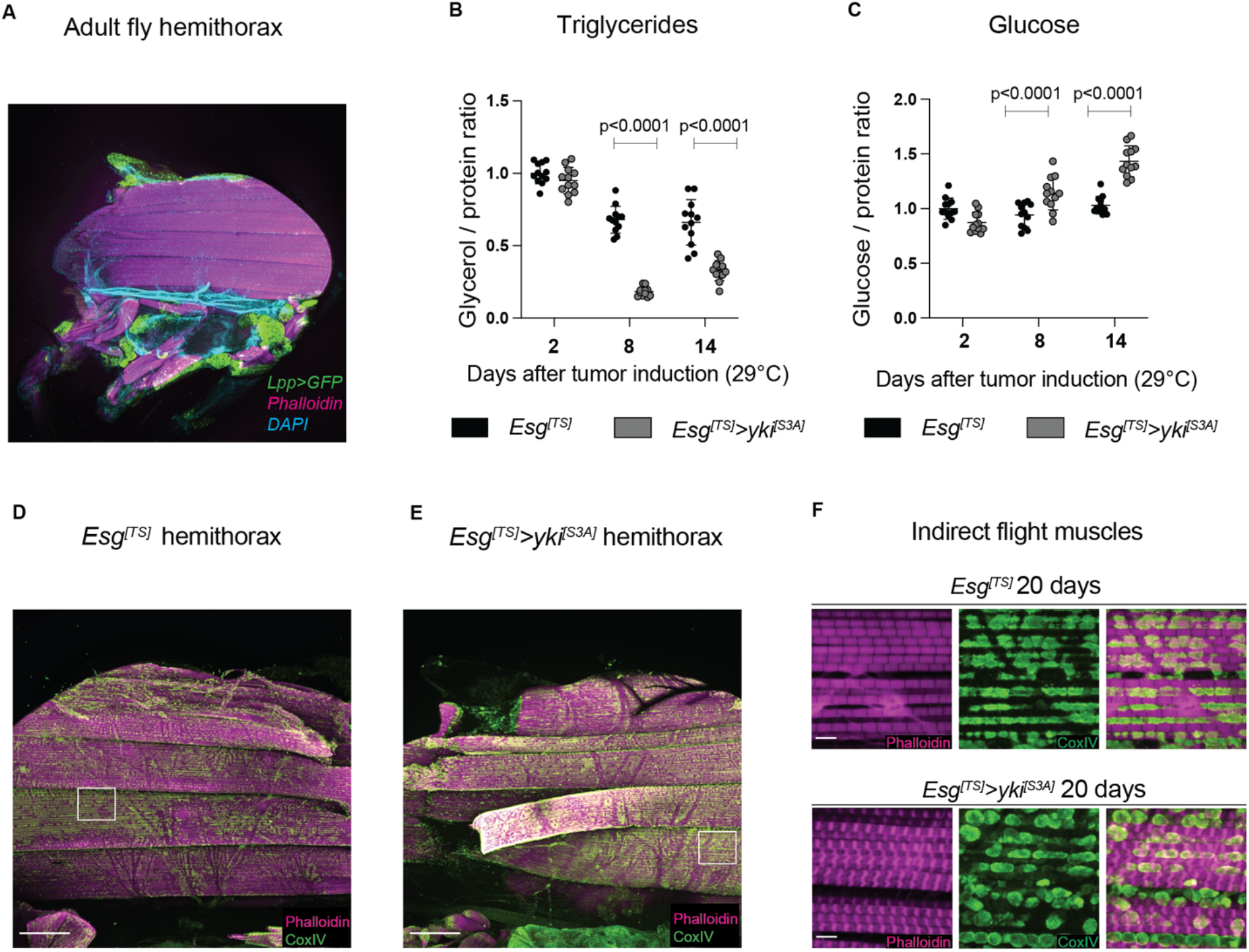
*Esg*^*[TS]*^>*yki*^*[S3A]*^ gut tumors induce systemic metabolic alterations and myofiber degradation. **(A)** *Lpp*^*[TS]*^*-GAL4* hemithorax stained with phalloidin (magenta) to mark the actin myofibers and expressing *UAS-TransTimer* (GFP only) to label fat body cells actively expressing *GAL4*. Trachea autofluorescence is detected in the DAPI channel. **(B, C)** TAG (B) and glucose content (C) in control *Esg*^*[TS]*^ or *Esg*^*[TS]*^*>yki*^*[S3A]*^ thoraces after tumor induction. TAG and glucose were calculated from the same homogenate. **(D-F)** Immunostaining of the flight muscles of control *Esg*^*[TS]*^ (D) or *Esg*^*[TS]*^*>yki*^*[S3A]*^ (E) hemithoraces after 20 days of tumor induction. Myofibrils are labelled with phalloidin (magenta) and mitochondria with CoxIV (green). Scale bar is 100 um. White boxes define magnified areas in (F). **(F)** Magnified areas of the flight muscles of control *Esg*^*[TS]*^ (D) or *Esg*^*[TS]*^*>yki*^*[S3A]*^ (E) hemithoraces scanned with a 60x objective lens. Sarcomere structure and mitochondrial morphology are severely altered in *Esg*^*[TS]*^*>yki*^*[S3A]*^ thoraces. Scale bar is 5 um. Values were normalized to the mean of control samples of 2 days after tumor induction (B, C). Statistical analysis was done using two-way ANOVA (B, C) with Sidak correction test for multiple comparisons (p<0.05). Data shows mean with SD.

Glucose is the main energy substrate used by flight muscles of *Drosophila*, whereas lipids appear to be unable to fuel continuous flight due to their slow processing(*20*). Insects that use carbohydrates as energy source to power flight muscles rely on aerobic glycolysis and oxidative metabolism to convert glucose to pyruvate for the TCA cycle(*21*). However, transcriptomic analysis of thoraces of flies with gut *yki-*tumors show a reduction in oxidative metabolism and glycolysis transcriptional profile(*14*), and flight muscles exhibit loss of function linked to wing positioning defects and mitochondrial dysfunction*(14)*. We therefore analyzed the thoracic dorso-longitudinal muscles (DLM), a subgroup of indirect flight muscles from flies with *yki-*tumors. Immunostaining showed myofiber degradation and compromised sarcomere structure (Fig. 1D-F, Fig. S1A-D). The percentage of *Esg*^*[TS]*^*>yki*^*[S3A]*^ flies with flight muscle degradation increased between 14 and 20 days after tumor induction, with most thoraces exhibiting myofiber degradation by 20 days (Fig. S1E, Table S1). This correlated with protein content in *Esg*^*[TS]*^*>yki*^*[S3A]*^ thoraces being significantly decreased between 14 and 20 days (Fig. S1F). In addition, degraded myofibers of *Esg*^*[TS]*^*>yki*^*[S3A]*^ thoraces had mitochondria with more circular morphology than the elongated mitochondria shown in control samples (Fig. 1D-F), suggesting mitochondrial dysfunction*(14)*. Thus, gut *yki*-tumors lead to changes in energy metabolism that precede signs of muscle degradation and resembles muscle wasting in cancer patients(*12, 13*).

In *Esg*^*[TS]*^*>yki*^*[S3A]*^ thoraces, p-AKT levels are reduced, an indicator of lower insulin signaling(*14*)(Fig. S2A). Further, given the negative regulation of the transcription factor FoxO by AKT(*22*), the expression of the FoxO target gene *4E-BP*(*22*) was elevated in *Esg*^*[TS]*^*>yki*^*[S3A]*^ thoraces(*14*) (Fig. S2B). Consistent with previous studies, suppression of *ImpL2* expression in *yki-* tumors restored *4E-BP* expression(*14*) (Fig. S2V). TAG reduction and glucose increase in thoraces was also ameliorated (Fig. S 2D,E), and the percentage of thoraces showing myofiber degradation was lower (Table S1, Fig. S2F). Given these results, we tested whether a muscle-specific reduction in insulin signaling would be sufficient to recapitulate the changes in TAG and glucose content and the myofiber degradation observed in *Esg*^*[TS]*^*>yki*^*[S3A]*^ thoraces. Forced expression of the p60 adaptor subunit, which mimics lower insulin signaling by reducing Pi3K activity(*23*), led to increased TAG content, while slightly reducing glucose levels in thoraces (Fig. S2H, I), opposite of what was observed in *Esg*^*[TS]*^*>yki*^*[S3A]*^ thoraces. Furthermore, p60 forced expression in muscles did not promote myofiber degradation even after 20 days of induction (Table S1). Interestingly, even though FoxO upregulation in muscles increases proteolysis in flight muscles(*24*), only 30% of thoraces showed myofiber degradation (Table S1). In addition, *FoxO* forced expression in muscles reduces TAG content in thoraces(*25*)(Fig. S2J), but glucose content only marginally increased, even after 20 days of sustained expression (Fig. S2K). Taken together, reducing insulin signaling specifically in muscle is not sufficient to drive either the metabolic changes or the myofiber degradation observed in *Esg*^*[TS]*^*>yki*^*[S3A]*^ thoraces, indicating that tumor-derived ImpL2 might be acting through additional mechanisms.

To identify novel genes responsible for modulating muscle metabolism in the context of gut *yki-*tumors, we performed a time-course RNA-seq experiment in thoraces spanning a tumor induction phase and a tumor “shutdown” phase by taking advantage of a GAL80 temperature-sensitive transgene (*tub-GAL80*^*[TS]*^) (Fig. S3A). We analyzed gene expression at day 14, a time point near the onset of myofiber degradation in which glucose content is higher in *Esg*^*[TS]*^*>yki*^*[S3A]*^ thoraces. During the tumor “shutdown” phase, which was initiated after 14 days of tumor induction and lasted 24 days until day 38 (Fig. S3A), TAG and glucose were restored to control levels, whereas the reduction in protein content was halted, indicating that the systemic effects of the *yki-* tumors were ameliorated (Fig. S3B-D). Also, the expression of both *yki*^*[S3A]*^ and *ImpL2* in the gut, and *4E-BP* in thoraces was restored (Fig. S3E-G), and only 50% of *Esg*^*[TS]*^*>yki*^*[S3A]*^ thoraces showed myofiber degradation (Table S1). Our analysis revealed 241 genes downregulated and 269 genes upregulated during the tumor induction phase, whose expression was then reversed during the “shutdown” phase (Table S2). GO-term enrichment and KEGG pathway analysis showed decreased expression of groups of genes related to mitochondria, glycolysis, generation of precursor metabolites and energy, and carbohydrate metabolism during the tumor induction phase (Table S3). Some of the increased groups included stress response genes, signal transduction and DNA-transcription factor activity (Table S3). Within this last category, we identified 23 transcription factors that were upregulated during tumor induction phase and could be candidate regulators of muscle metabolism (Fig. S3H). We focused on *REPTOR*, a transcription factor regulated by TOR signaling, previously linked to the regulation of metabolism in *Drosophila* larvae(*26*). The expression of *REPTOR* was increased in *Esg*^*[TS]*^*>yki*^*[S3A]*^ thoraces (Fig. 2A), restored during the tumor “shutdown” phase (Fig S3I), and tumor-derived *ImpL2* could modulate *REPTOR* transcription in thoraces (Fig. 2B).

**Fig. 2:**
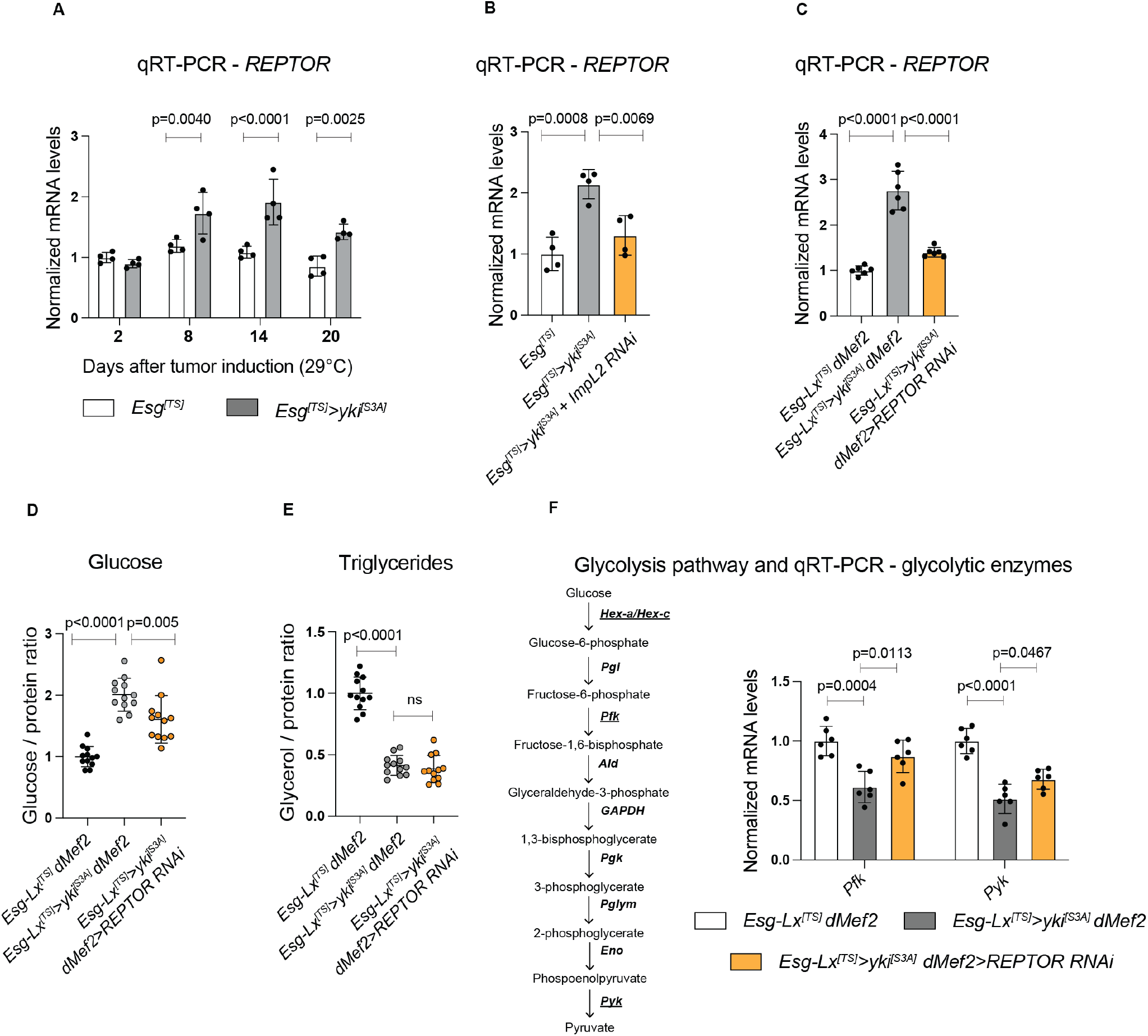
REPTOR activity in muscle regulates glucose content and glycolysis in *Esg*^*[TS]*^>*yki*^*[S3A]*^ thoraces. **(A-C)** *REPTOR* mRNA levels in: (A) control *Esg*^*[TS]*^ and *Esg*^*[TS]*^*>yki*^*[S3A]*^ thoraces after tumor induction; (B) in *Esg*^*[TS]*^*>yki*^*[S3A]*^ thoraces with *ImpL2* knockdown in the gut *yki-*tumors after 20 days of tumor induction; (C) in thoraces with simultaneous induction of gut *yki-*tumors (*Esg*-LexA/LexAop-*yki*^*[S3A]*^) and muscle-specific *(dMef2-GAL4)* knockdown of *REPTOR* RNAi after 12 days of tumor induction. *REPTOR* is upregulated in *Esg*^*[TS]*^*>yki*^*[S3A]*^ thoraces and ImpL2 modulates *REPTOR* expression. Knockdown of *REPTOR* in muscles significantly reduces its expression in *Esg*^*[TS]*^*>yki*^*[S3A]*^ thoraces. **(D, E)** Glucose (D) or TAG (E) content in *Esg*^*[TS]*^*>yki*^*[S3A]*^ thoraces when *REPTOR* expression is reduced in muscles after 12 days of tumor induction. TAG and glucose were calculated from the same homogenate. **(F)** Gene expression analysis of the glycolytic rate-limiting enzymes *pfk* and *pyk* after 12 days of tumor induction. Values were normalized to the mean of control samples of 2 days after tumor induction (A) or mean of the control samples (B-F). Statistical analysis was done using two-way ANOVA (A) or one-way ANOVA (B-F) with Sidak correction test for multiple comparisons (p<0.05). Data shows mean with SD.

REPTOR activity has been previously shown to be modulated through phosphorylation by the mTOR pathway in larvae, leading to its cytoplasmic retention(*26*). Even though insulin signaling was reduced in *Esg*^*[TS]*^*>yki*^*[S3A]*^ thoraces, we could not detect significant changes in the levels of phosphorylation of the direct mTOR target S6K (Fig. S4A), suggesting that mTOR activity was not decreased in *Esg*^*[TS]*^*>yki*^*[S3A]*^ thoraces. Nevertheless, the elevated *REPTOR* mRNA indicated that it could be more active in muscles of *Esg*^*[TS]*^*>yki*^*[S3A]*^ thoraces. We used a second binary system of gene expression, the *LexA-LexAop* system(*27*), to induce gut *yki-*tumors, while independently decreasing *REPTOR* expression in muscle using *dMef2*^*[TS]*^*-GAL4* REPTOR*-RNAi*. Knockdown of *REPTOR* in muscles of *Esg*^*[TS]*^*>yki*^*[S3A]*^ thoraces reduced the elevation of its transcript (Fig. 2C) without affecting tumor growth (Fig. S4B-D). However, we could not retrieve flies after 12 days of tumor induction that maintained a strong suppression of *REPTOR* expression in muscles, suggesting that the upregulation of *REPTOR* might be necessary for survival of flies with *yki-*tumors. Importantly, the increase in glucose content observed in *Esg*^*[TS]*^*>yki*^*[S3A]*^ thoraces was significantly blunted after 12 days of tumor induction (Fig. 2D), showing that *REPTOR* upregulation in muscles of *Esg*^*[TS]*^*>yki*^*[S3A]*^ flies was necessary to modulate glucose levels in thoraces. Also, knockdown of *REPTOR* in wildtype muscles did not affect TAG or glucose levels (Fig. S4E,F), indicating that the effect of REPTOR on glucose content was specific to *Esg*^*[TS]*^*>yki*^*[S3A]*^ flies. Thus, an increase in REPTOR activity in muscles regulates glucose metabolism in the context of *yki-*tumors.

Transcriptomic analysis of *Esg*^*[TS]*^*>yki*^*[S3A]*^ thoraces showed a severe reduction in energy metabolism gene signatures, in particular glycolysis(*14*) (Table S3). We therefore considered whether higher levels of REPTOR in muscles could be modulating the expression of rate-limiting glycolytic enzymes in this tumor context. Whereas we could not detect significant changes in the expression of *Hexokinase-a* and *Hexokinase-*c in *Esg*^*[TS]*^*>yki*^*[S3A]*^ thoraces (Fig. S4G), both *Phosphofructokinase (Pfk)* and *Pyruvate kinase (Pyk)* were downregulated (Fig. 2F). Importantly, reduction of *REPTOR* mRNA levels in *Esg*^*[TS]*^*>yki*^*[S3A]*^ muscles fully restored the expression of *pfk* to control levels, and partially rescued *pyk* expression (Fig. 2F), suggesting that *REPTOR* contributes to the reduction in glycolytic gene expression in muscle during tumor progression. Given the role of REPTOR in modulating *pfk* expression in the context of *yki-*tumors, we tested whether REPTOR could have a similar role under nutritional restriction, a setting in which mitochondria preferentially use lipids instead of glucose. *REPTOR* mRNA levels were increased in muscles after a short period of starvation (Fig. S4H). Notably, *pfk* expression in starved flies was increased when *REPTOR* was suppressed in muscles, thus demonstrating that REPTOR represses expression of *pfk* either in the presence of *yki-*tumors or under nutritional restriction (Fig. S4I). Thus, our results suggest that an increase in *REPTOR* activity in muscles might be a response to gut *yki-*tumors to suppress the use of glucose as an energy substrate in peripheral tissues.

Increased levels of insulin signaling in muscles of *yki-*flies has recently been shown to reduce the percentage of flies with wing position defects and to rescue systemic organ wasting(*28*). In *Drosophila* larval muscles, insulin signaling induces glycolysis to regulate growth and myoblast fusion(*29*), suggesting that the increase in insulin signaling in muscles of *yki-*flies might have a protective effect by increasing glycolysis in muscles. Forced expression of an active form of *AKT* in muscle (*dMef2*^*[TS]*^*>Myr-AKT*) led to an amelioration of myofiber degradation and to suppression of *4E-BP* upregulation (Fig. S5A,B, Table S1) without affecting tumor growth (Fig. S5B-D). This rescue correlated with an increase in expression of *pyk* and *pfk* in *Esg*^*[TS]*^*>yki*^*[S3A]*^ *dMef2*^*[TS]*^*>Myr-AKT* thoraces (Fig. 2H). Because the adult thorax is mainly composed of muscle fibers, we quantified ATP content as proxy for changes in muscle energy metabolism. Control and *Esg*^*[TS]*^*>yki*^*[S3A]*^ thoraces had similar levels of ATP at 12 days after tumor induction, but by day 16 ATP content was significantly decreased in *Esg*^*[TS]*^*>yki*^*[S3A]*^ thoraces. *Myr-AKT* expression significantly increased ATP content in *Esg*^*[TS]*^*>yki*^*[S3A]*^ thoraces at both time points, indicating a rescue in ATP content, although it did not completely prevent the gradual decline of ATP levels over time caused by *yki*-tumors (Fig. 2G). The protective effects of *Myr-AKT* were not likely caused by a change in expression or activity of REPTOR, since both the transcript level and the amount of p-S6K, a readout of mTOR pathway activity known to modulate REPTOR(*26*), were unaltered (Fig. S5E,F). Altogether, our results suggest that alterations of energy metabolism and myofiber degradation in *Esg*^*[TS]*^*>yki*^*[S3A]*^ thoraces may, at least in part, be caused by changes in glucose metabolism by increased REPTOR activity and decreased insulin signaling in muscles.

*Yki-*tumors produce multiple factors that can induce complex systemic metabolic changes(*14, 30, 31*), making it difficult to isolate the effects of REPTOR in muscle metabolism. To study *REPTOR* in normal muscles in the absence of gut *yki-*tumors, we overexpressed in muscles (*dMef2*^*[TS]*^*-GAL4*) a wildtype allele *REPTOR*^*[WT]*^ or a constitutively active form of *REPTOR, REPTOR*^*[ACT]*^, that cannot be phosphorylated by TORC1(*26*). Both genes upregulated *4E-BP* expression(*26*) (Fig. S6A), but only *REPTOR*^*[ACT]*^ was sufficient to increase glucose content in thoraces after 4 days of expression (Fig 3A, Fig. S6B). Almost 100% of thoraces displayed myofiber degradation after 8 days of expression (Table S1), with degraded myofibers showing altered mitochondrial morphology (Fig 3B). In contrast, sustained overexpression in muscles of *REPTOR*^*[WT]*^, or reducing the activity of the mTOR pathway, which controls the activity of REPTOR(*26*), did not affect glucose content or promoted myofiber degradation (Table S1, Fig. S6B,C). Thus, an increase in the activity of REPTOR in wildtype muscles is sufficient to recapitulate the glucose content increase and the myofiber degradation phenotypes induced by *yki-* tumors.

**Fig. 3:**
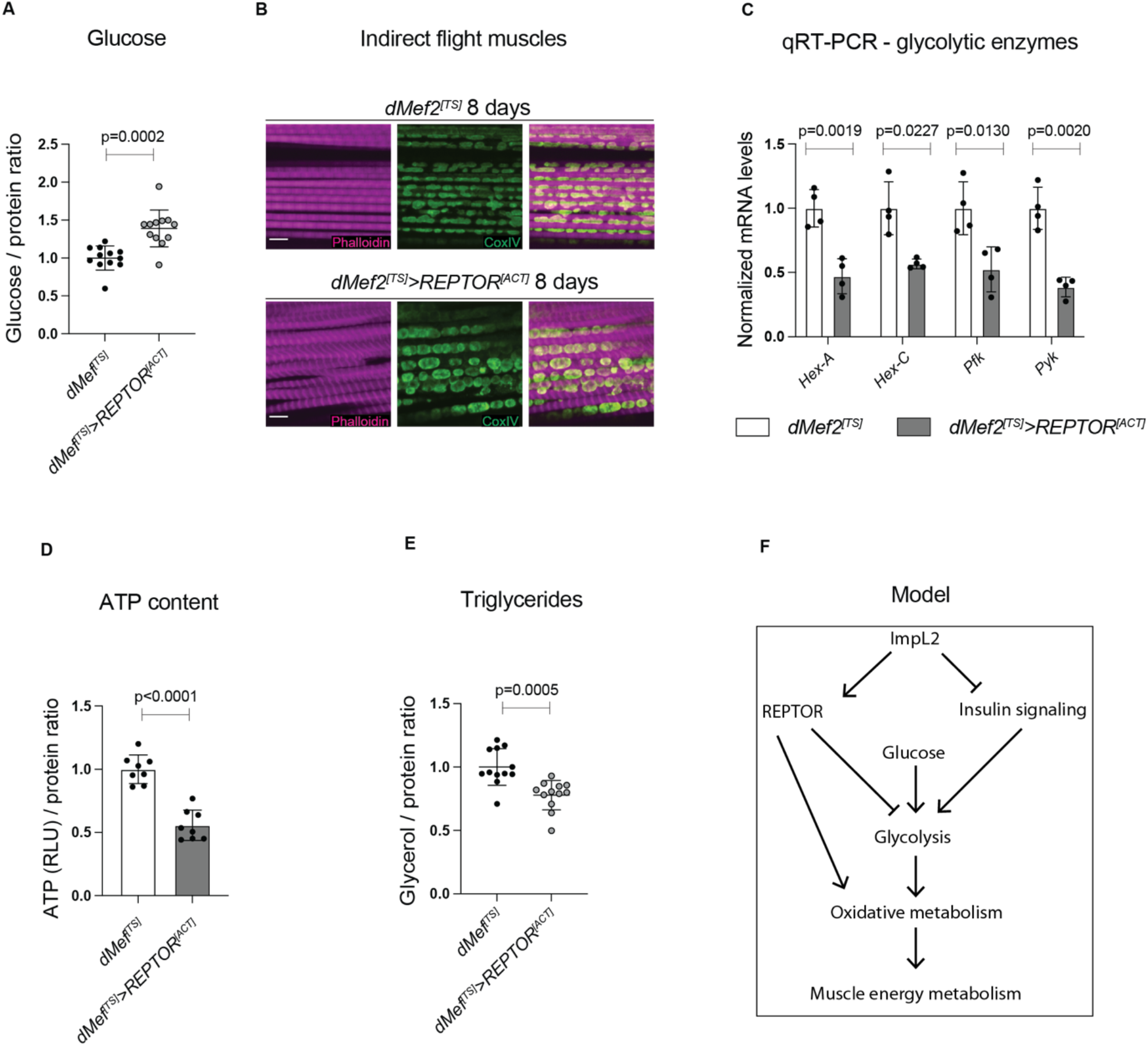
REPTOR overexpression in muscle modulates glucose metabolism and represses glycolysis. **(A)** Glucose content in thoraces when an active allele of *REPTOR* (*REPTOR*^*[ACT]*^) is overexpressed in muscles (*dMef2-GAL4*) for 4 days. **(B)** Immunostaining of the flight muscles of hemithoraces after 8 days of *REPTOR*^*[ACT]*^ induction in muscles. Myofibrils are labelled with phalloidin (magenta) and mitochondria with CoxIV (green). Scale bar is 5 um. **(C)** Gene expression analysis of the four glycolytic rate-limiting enzymes (*hex-a, hex-c, pfk* and *pyk*) after 4 days of overexpression of *REPTOR*^*[ACT]*^ in muscles. **(D)** ATP content in thoraces after 4 days of expression. Values were pooled from two independent experiments. **(E)** TAG content in thoraces when *REPTOR*^*[ACT]*^ is overexpressed in muscles for 4 days. TAG was calculated from the same samples as in (A). **(F)** Model of the role of REPTOR in muscles. ImpL2 positively regulates REPTOR while reducing insulin signaling in muscles. Glucose usage is regulated by the action of REPTOR and insulin signaling. REPTOR potentially promotes oxidative phosphorylation. All values were normalized to the mean of control samples (A, C, D, E). Statistical analysis was done using two-tailed *t*-test with Welch’s correction (A, C, D, E, p<0.05). Data shows mean with SD.

To further understand the role of REPTOR in energy metabolism we generated stable S2R+ cell lines with CuSO4-inducible *REPTOR*^*[WT]*^ or *REPTOR*^*[ACT]*^ alleles to test whether REPTOR controlled glycolysis and oxidative metabolism rates in a cell-autonomous manner. Even though both transgenes showed transcriptional activity, *REPTOR*^*[ACT]*^ exhibited higher target gene induction than *REPTOR*^*[WT]*^ (Fig. S6D). We then measured oxygen consumption and extracellular acid production in these S2R+ cell lines. From these values, the relative contributions of mitochondrial respiration and glycolysis to overall cellular ATP production can be derived(*32*). Notably, only S2R+ cells expressing *REPTOR*^*[ACT]*^ displayed a significant increase in mitochondrial respiration, though without changes in glycolytic contribution to ATP production (Fig. S6E). Nevertheless, intracellular lactate in S2R+ cells expressing *REPTOR*^*[ACT]*^ was reduced, consistent with an increase in oxygen consumption and mitochondrial respiration (Fig. S6F,G).

Next, we tested *in vivo* for effects of REPTOR on glycolysis and oxidative phosphorylation. Gene set enrichment analysis of thoraces with sustained expression of *REPTOR*^*[ACT]*^ in muscles showed a downregulation of genes involved in energy metabolism, in particular, glycolysis, but also oxidative phosphorylation and mitochondria (Table S4, 5), indicating an impairment of energy metabolism similar to that observed in the transcriptional profile of the *Esg*^*[TS]*^*>yki*^*[S3A]*^ thoraces(*14*) (Table S3). Consistent with this, *REPTOR*^*[ACT]*^ significantly reduced the expression of glycolysis rate-limiting enzymes in thoraces (Fig 3C), as well as ATP content (Fig. 3D). Even though these results contrast with the lack of effect of REPTOR in decreasing the contribution of glycolysis for ATP production in S2R+ cells (Fig. S6E), TAG content in thoraces was reduced suggesting that prolonged *REPTOR*^*[ACT]*^ overexpression in muscles may promote a shift to lipids as an energy substrate (Fig. 3E). Yet, these observations indicate that unlike in the *in vitro* system of S2R+ cells, sustained *REPTOR* overexpression *in vivo* in muscle may lead to the collapse of the oxidative phosphorylation program, possibly due to a combination of slow metabolism of lipids in flight muscles and their reliance on glucose(*20*). Nevertheless, our results both *in vivo* and *in vitro* suggest that *REPTOR* works as a regulator of metabolic flexibility in muscle, likely by reducing glucose metabolism and glycolysis while promoting oxidative phosphorylation (Fig. 3F).

To assess the evolutionary conservation of REPTOR function in the regulation of muscle metabolism, we examined CREBRF, the mammalian ortholog of REPTOR. The human CREBRF protein shares 16% identity and 25% similarity with REPTOR, although the REPTOR residues phosphorylated by mTOR (S527 and S530) do not appear to be conserved in CREBRF. Nevertheless, *Crebrf* expression has been reported to increase upon mTOR inhibition, and its forced expression protects against starvation in cultured adipocytes, suggesting that its activity may be responsive to nutrient availability(*33, 34*). Moreover, a *Crebrf* coding polymorphism associated with obesity in the Samoan population has the strongest effect on BMI of any common obesity-risk variant(*34*).

We first examined the regulation of the endogenous *Crebrf* gene. In mice, the *Crebrf* transcript was increased in muscle upon fasting and suppressed upon refeeding, confirming its regulation by nutritional availability (Fig. 4A). Similarly, in mouse myotubes *in vitro, Crebrf* was increased upon the stress of nutrient withdrawal (Fig. 4B), whereas it was decreased by insulin signaling (Fig. 4C).

**Fig. 4.**
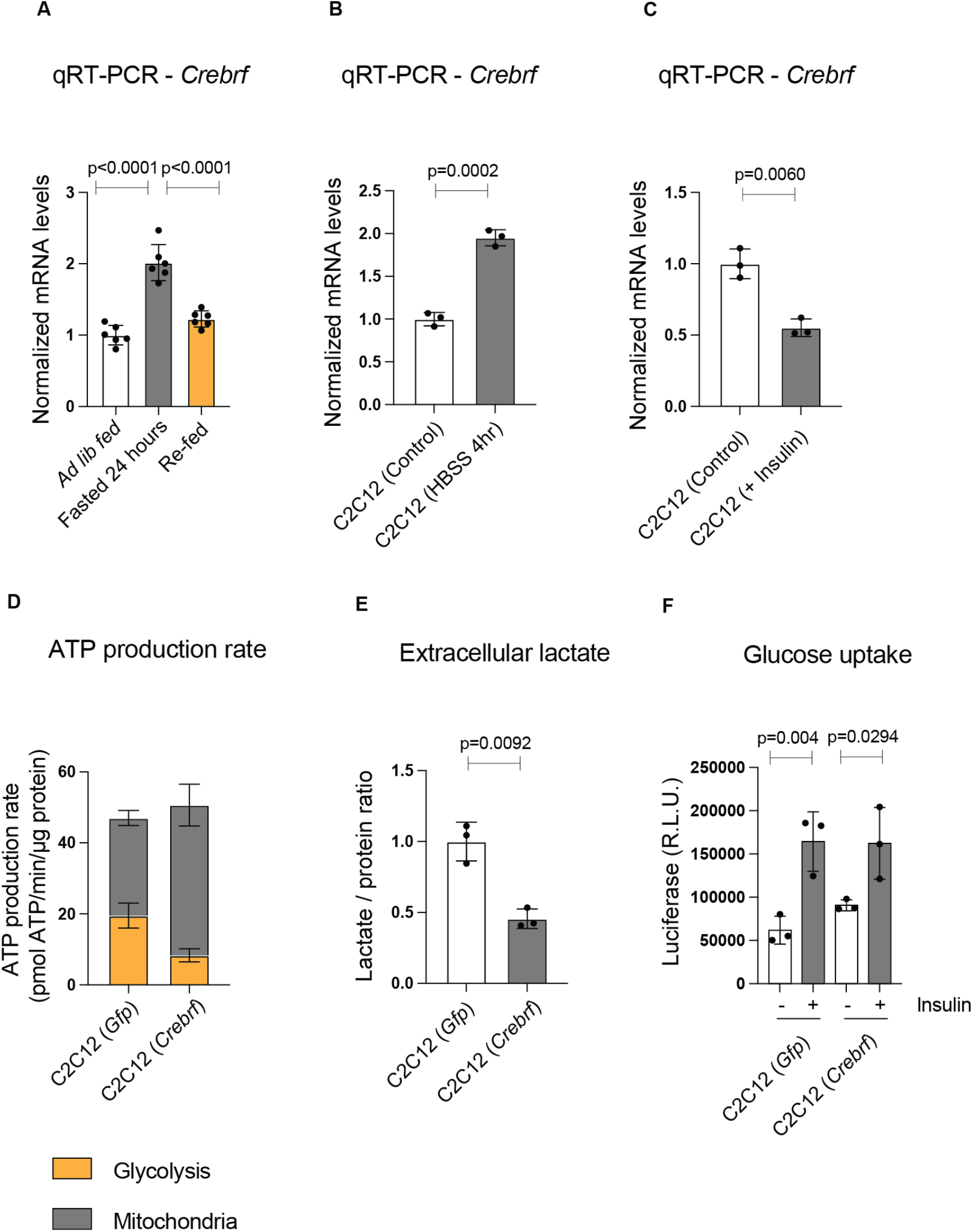
CREBRF is a repressor of glycolysis in mammalian myotubes. **(A-C)** *Crebrf* mRNA levels in mouse quadriceps muscle upon *ad libitum* feeding, 24 hr fast or 4 hr re-feeding (A), in C2C12 myotubes incubated in HBSS for 4 hr (B) or treated with insulin (100 nM) (C). *Crebrf* levels in muscle are modulated *in vivo* and *in vitro* by the nutritional status. **(D)** ATP production rate contributed by glycolysis and mitochondrial respiration in C2C12 myotubes upon adenoviral expression of *Crebrf*. CREBRF represses glycolysis while increasing mitochondrial contribution to ATP production. Values for glycolysis and mitochondrial respiration were calculated from ten individual wells for each condition. **(E)** Secreted lactate measurement in media conditioned by C2C12 myotubes expressing *Crebrf*. Sustained expression of *Crebrf* reduces lactate levels in the media, indicating lower glycolysis rate. **(F)** Basal and insulin-stimulated glucose uptake by myotubes expressing *Crebrf*. The role of CREBRF in repressing glycolysis is not due to decreased glucose uptake by myotubes. All values were normalized to the average of the control samples (A-C, E, F). Statistical analysis was done using one-way ANOVA (A) with Sidak correction test for multiple comparisons (p<0.05) or two-tailed *t*-test with Welch’s correction (B, C, E, F). Data shows mean with SD.

Next, we examined the effects of forced *Crebrf* expression on energy metabolism in myotubes. We transduced C2C12 myotubes with adenovirus encoding *Crebrf* and measured the relative contributions of mitochondrial respiration and glycolysis to overall cellular ATP production. In control cells, glycolysis and mitochondrial respiration each contributed approximately half of the cellular ATP production. In contrast, cells expressing *Crebrf* showed a substantial shift away from glycolysis: their absolute rate of glycolytic ATP production decreased by 60%, alongside a concomitant 40% increase in the absolute rate of mitochondrial ATP production (Fig. 4D). This effect was independent of the energy substrates available to myotubes, as it was observed not only in the presence of glucose, pyruvate, and glutamine but also in media lacking all three of these components (Fig. S7A). As expected, lactate production was halved in cells expressing *Crebrf*, mirroring their reduced glycolytic ATP production (Fig. 4E). Cells expressing *Crebrf* exhibited unaltered glucose uptake rates in basal as well as in insulin-stimulated conditions (Fig. 4F), suggesting that their reduced glycolytic flux was caused by a reduction in the intracellular steps of glycolysis.

Given the role of CREBRF in transcriptional regulation, we next used RNA sequencing to assess the gene expression changes associated with the altered energy metabolism of myotubes expressing *Crebrf*. Unbiased gene set enrichment analysis revealed oxidative phosphorylation metabolism as the top upregulated term, whereas the top downregulated terms were interferon response, TNFα signaling, and hypoxia (Fig. S7B). Moreover, the glycolysis gene set itself was significantly decreased (Fig. S7C). The concerted upregulation of mRNAs encoding many mitochondrial proteins suggested a change in activity of an upstream regulator. Indeed, the protein levels of PGC1α, a transcriptional co-activator sufficient to drive mitochondrial biogenesis, were increased in cells expressing *Crebrf*, as were mRNA levels of known PGC1α target genes (Fig. S7D,E, Table S6). Together, these results suggest that *Crebrf* drives gene expression changes that elevate mitochondrial respiration and decrease glycolysis in myotubes.

Our results demonstrate that REPTOR and CREBRF promote oxidative metabolism *in vitro*. In flies, higher activity of *REPTOR* in muscle increased glucose content and reduced *pfk* expression, likely reflecting reduced glucose utilization, while also decreasing TAG content. In mammalian myotubes, *Crebrf* gain-of-function also repressed glycolysis, resulting in a shift towards mitochondrial respiration that was accompanied by broad increases in the oxidative phosphorylation gene expression program. The fact that both REPTOR and CREBRF can promote oxidative metabolism suggests that these orthologs are involved in a metabolic response to unfavorable nutrient availability, as observed in the context of *yki*-tumors or nutritional restriction. Increased REPTOR activity in *Drosophila* muscles might function, in the context of *yki*-tumors, as a systemic response that limits glucose use in peripheral tissues. However, sustained activation of REPTOR in muscle eventually led to a collapse of both the glycolysis and oxidative phosphorylation gene expression programs, as well as alterations of mitochondrial morphology and reduced ATP content. This may reflect the dependence of *Drosophila* flight muscles on glucose as the preferred energy source(*20*). Together, our work implicates REPTOR and CREBRF as powerful modulators of metabolic flexibility and energy substrate choice in muscle.

## Acknowledgments

We are grateful to David Doupe and Jonathan Zirin for comments on the manuscript, Justin Bosch and Nirmalya Chatterjee for sharing plasmids, Ben Ewen-Campen for sharing qPCR primers, Aurelio Teleman for providing the REPTOR reagents, Raghuvir Viswanatha for help with tissue culture, the TRiP at Harvard Medical School, BDSC and NIG for providing transgenic fly stocks used in this study, Frank Schnorrer, Maria Spletter and Nuno Luis for advice on thorax dissections, Christians Villalta for fly embryo injections, Yuan Feng for technical help with dissections and RNA extractions, Jenny Ro for technical help with the colorimetric assays, Paula Montero and the MicroN Core Facility for assistance with confocal microscopy, the BPF Next-Gen Sequencing Core Facility at Harvard Medical School for the RNA-sequencing services. PS was supported by a Human Frontiers Science Program long-term Fellowship LT000937/2016. This work was supported by NIH R01AR057352 and P01CA120964 to NP, NIH grants R01DK123228 and R01 DK119117 to BMS and DRCRF Fellowship DRG-120-17 to PAD. NP is an investigator of HHMI.

## Author contributions

PS and NP designed the study in *Drosophila*. PS performed and analyzed the experiments with *Drosophila* and S2R+ cells. RB and EF provided technical help with *Drosophila* experiments. PS and PAD designed the experiments with mouse and C2C12 myotubes. PAD performed and analyzed the Seahorse assays and the experiments with mouse and C2C12 myotubes with technical help from SEW. YH, HW, JR and PAD analyzed the RNA-seq datasets. PJ generated the UAS-HA-FoxO reagents and performed the western blots for p-S6K. PS, PAD and NP wrote the manuscript. BMS and PJ provided critical feedback to the manuscript.

---

**Fig. S1:**
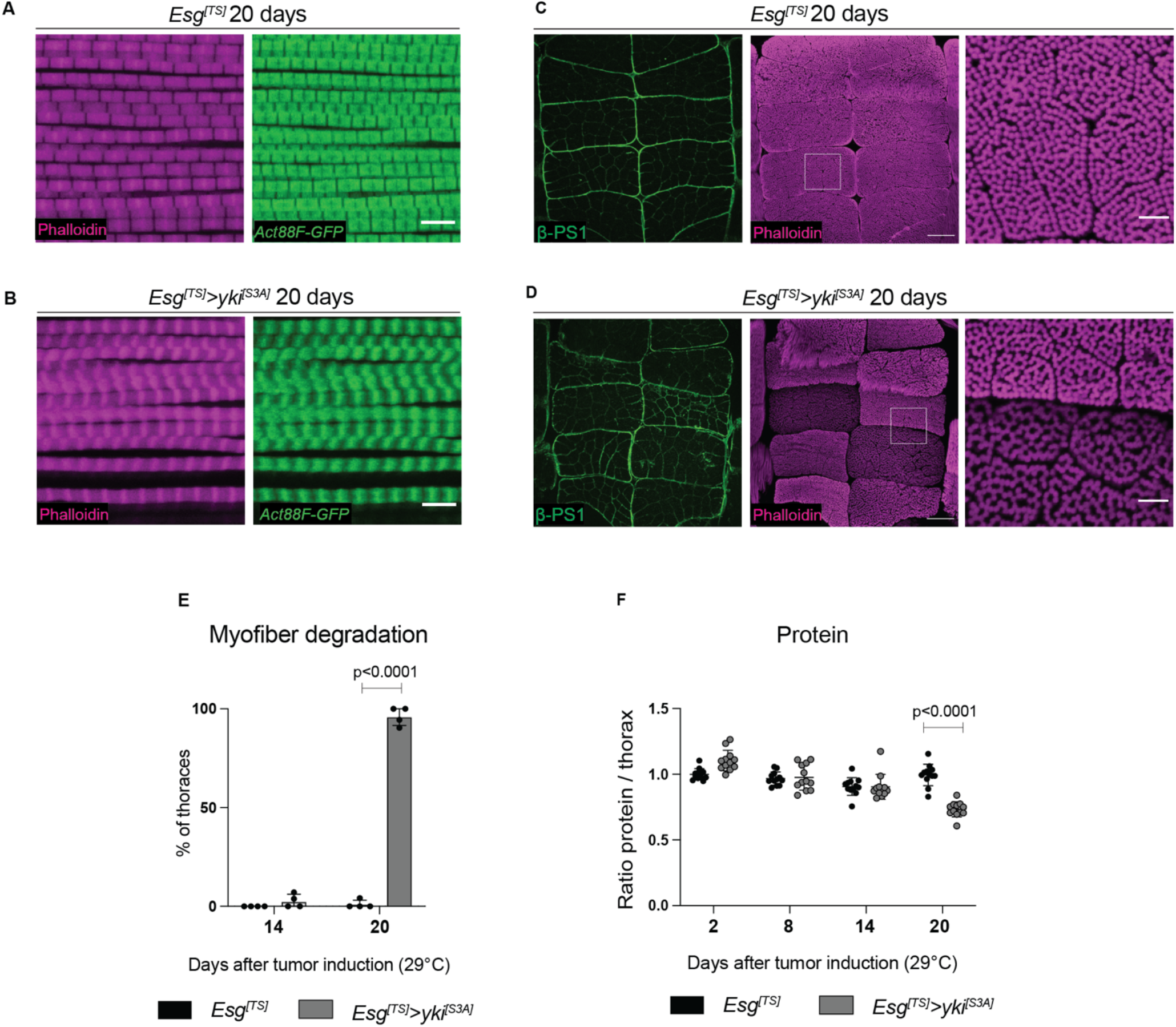
Characterization of the muscle degradation phenotype in *Esg*^*[TS]*^>*yki*^*[S3A]*^ thoraces. **(A, B)** Magnified areas of flight muscles expressing Actin88F::GFP (green) and stained with phalloidin (magenta) to label the sarcomeres in *Esg*^*[TS]*^ (A) or *Esg*^*[TS]*^*>yki*^*[S3A]*^ thoraces (B) after 20 days of tumor induction. Sarcomeres are shorter and myofiber structure is severely affected in *Esg*^*[TS]*^*>yki*^*[S3A]*^ thoraces. **(C, D)** Cross-section of control *Esg*^*[TS]*^ (A) or *Esg*^*[TS]*^*>yki*^*[S3A]*^ thoraces (B) after 20 days of tumor induction labeled with βPS1 (green) and phalloidin (magenta). Scale bar is 50 um. White boxes indicate zoomed areas in the right panels. Scale bar in magnified areas is 10 um. **(E)** Percentage of thoraces showing indirect flight muscles with at least one myofiber degraded after 14 or 20 days of tumor induction in the gut (Table S1). Myofiber degradation increases after 14 days of tumor induction. **(F)** Protein content in control *Esg*^*[TS]*^ (a) or *Esg*^*[TS]*^*>yki*^*[S3A]*^ thoraces. Protein content was calculated from the same samples as used in Fig. 1B,C. Each dot represents 8 thoraces. *Esg*^*[TS]*^*>yki*^*[S3A]*^ thoraces show lower protein content at 20 days after tumor induction. Values were normalized to the mean of control samples of 2 days after tumor induction (F). Statistical analysis was done using two-way ANOVA (E, F) with Sidak correction test for multiple comparisons (p<0.05). Data shows mean with SD.

**Fig. S2:**
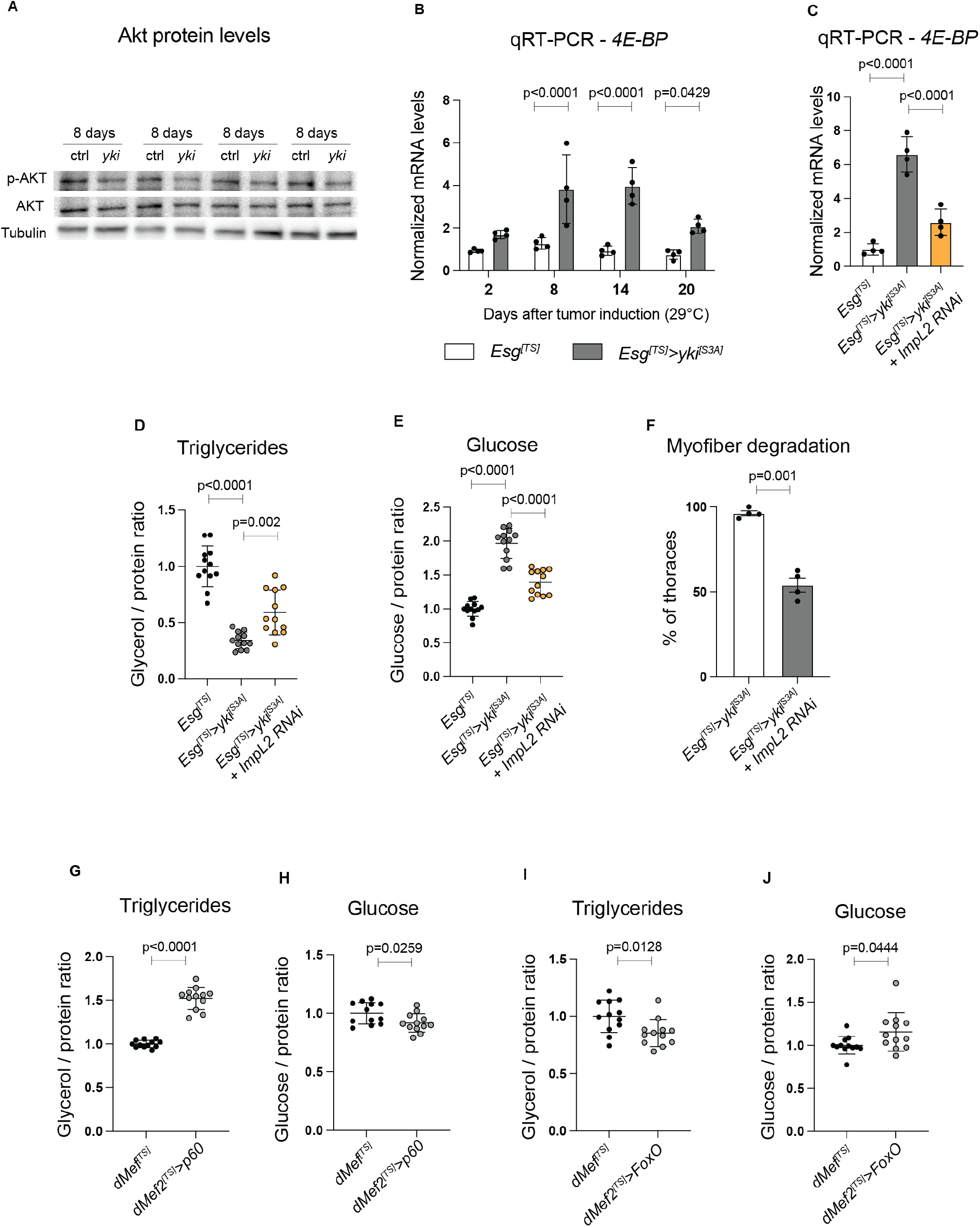
Insulin signaling reduction in muscles is not sufficient to promote the phenotypes observed in *Esg*^*[TS]*^>*yki*^*[S3A]*^. **(A)** Protein levels of p-AKT and total AKT from control *Esg*^*[TS]*^ or *Esg*^*[TS]*^*>yki*^*[S3A]*^ thoraces after 8 days of tumor induction. Four biological replicates are shown. p-AKT levels are lower in *Esg*^*[TS]*^*>yki*^*[S3A]*^ thoraces. **(B)** mRNA levels of *4E-BP Esg*^*[TS]*^ or *Esg*^*[TS]*^*>yki*^*[S3A]*^ thoraces after tumor induction. **(C-F)** Knockdown of *ImpL2* in gut *yki-*tumors. **(C)** mRNA levels of *4E-BP* in thoraces after 20 days of tumor induction. **(D, E)** TAG content in thoraces after 8 days of tumor induction (D) and glucose after 14 days of tumor induction (E). **(F)** Percentage of thoraces showing myofiber degradation after 20 days after tumor induction. See Table S1. Only *Esg*^*[TS]*^*>yki*^*[S3A]*^ and *Esg*^*[TS]*^*>yki*^*[S3A]*^ + ImpL2 RNAi are shown. (**G-J)** TAG and glucose content in thoraces when *p60* (G, H) or *FoxO* (I, J) are overexpressed in muscles. Glucose and TAG were quantified from the same samples. Overexpression of *p60* or *FoxO* in muscle does not induce the metabolic phenotypes observed in *Esg*^*[TS]*^*>yki*^*[S3A]*^ thoraces. Values were normalized to the mean of control samples of 2 days of tumor induction (B) or mean of the control samples (C-E, G-H). Statistical analysis was done using two-way ANOVA (B) or one-way ANOVA (C-E) with Sidak correction test for multiple comparisons (p<0.05) or using two-tailed *t*-test with Welch’s correction (F-J, p<0.05). Data shows mean with SD.

**Fig. S3:**
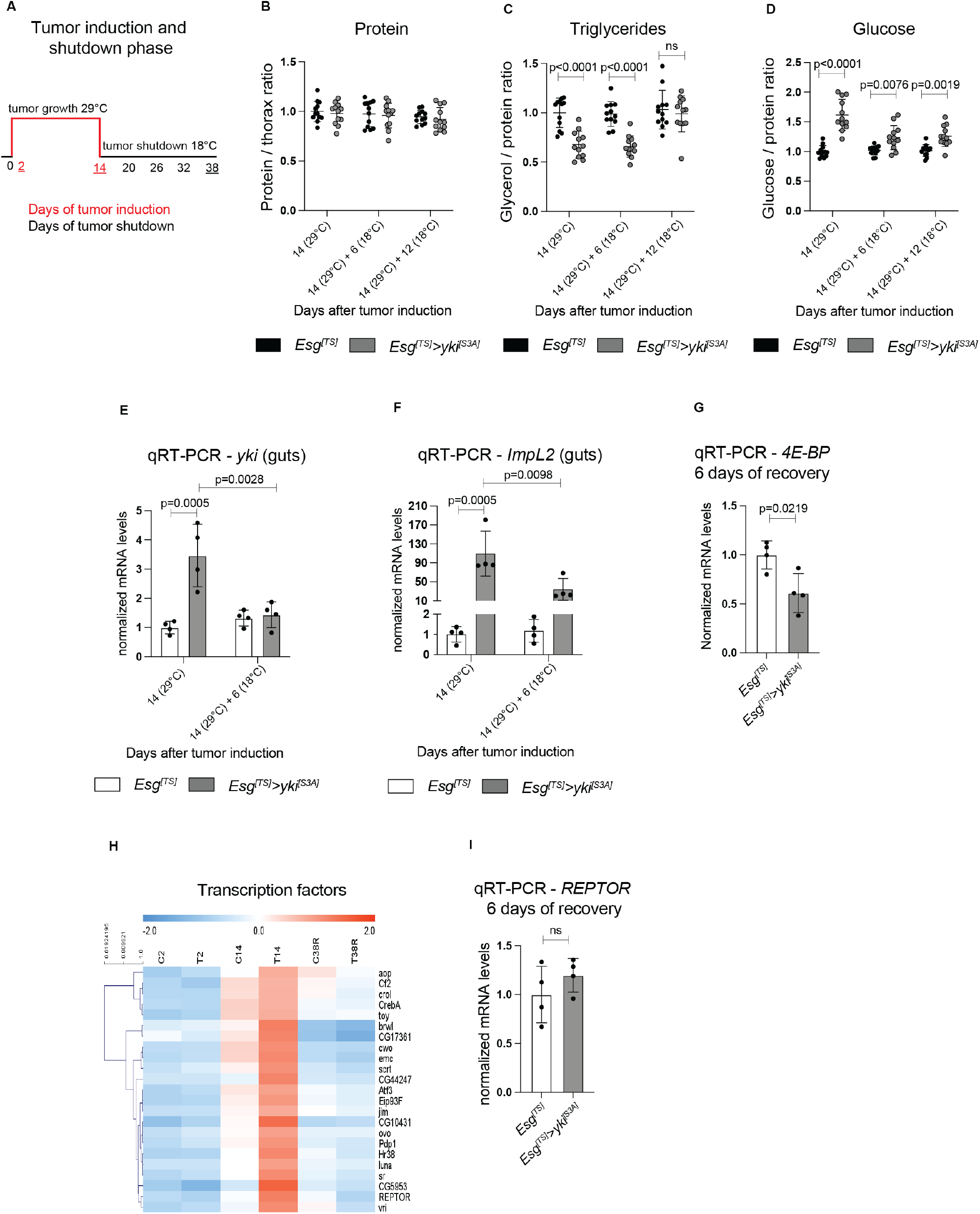
Time-course RNA-seq of *Esg*^*[TS]*^>*yki*^*[S3A]*^ thoraces. **(A)** Model showing the duration of the tumor induction (red line - 29°C) and the tumor shutdown phase (black line - 18°C). Underlined days were used as time points for RNA-seq analysis of *Esg*^*[TS]*^*>yki*^*[S3A]*^ thoraces. Three time points were selected for analysis, 2 and 14 days after tumor induction, and 24 days of tumor shutdown (38 days). **(B-D)** Protein (B), TAG (C) and glucose (D) content in thoraces during the recovery phase after 14 days of tumor induction. Values were calculated from the same samples. Each dot represents 8 thoraces. **(E, F)** mRNA levels of *yki* (E) and *ImpL2* (F) in control and *Esg*^*[TS]*^*>yki*^*[S3A]*^ guts after 14 days of tumor induction and after 6 days of recovery. *Yki* and *ImpL2* expression is significantly reduced after 6 days at 18°C. **(G)** mRNA levels of *4E-BP* in control and *Esg*^*[TS]*^*>yki*^*[S3A]*^ thoraces after 6 days of recovery at 18°C. Values were normalized to *rp49*. **(H)** Heat map showing the transcription factors with reverse pattern of expression for control (C) and *Esg*^*[TS]*^*>yki*^*[S3A]*^ (T) thoraces during tumor induction (C2 to C14; T2 to T14) and recovery phase (C38R and T38R). **(I)** mRNA levels of *REPTOR* in *Esg*^*[TS]*^*>yki*^*[S3A]*^ thoraces after 6 days of recovery. *REPTOR* levels are restored after 6 days of tumor shutdown (18°C). Values were normalized to *rp49*. Statistical analysis was done by two-way ANOVA (B-F) with Sidak multiple comparison correction (p<0.05) or using two-tailed *t*-test with Welch’s correction (G, I, p<0.05). Data shows mean with SD.

**Fig. S4:**
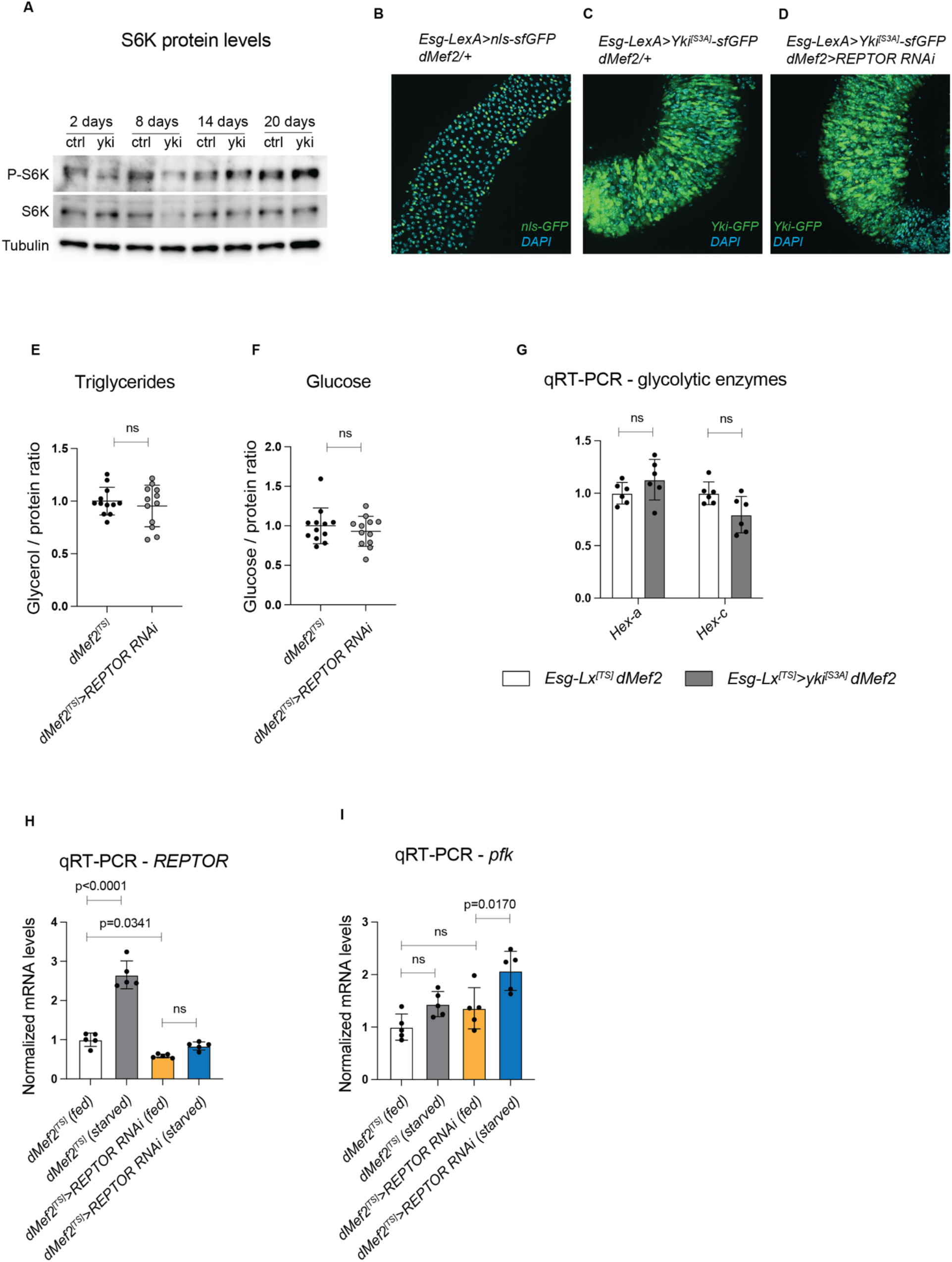
Characterization of the metabolic role of REPTOR in muscle in the context of *yki-*tumors or nutritional restriction. **(A)** Protein levels of p-S6K and total S6K from control (*Esg*^*[TS]*^) or *yki* (*Esg*^*[TS]*^*>yki*^*[S3A]*^) thoraces. No significant changes in p-S6K are detected in *Esg*^*[TS]*^*>yki*^*[S3A]*^ thoraces. **(B-D)** Immunostaining of posterior midguts after 12 days of tumor induction with the *esg-*LexA/LexAop-*yki*^*[S3A]*^ (*EsgLx*^*[TS]*^*>yki*^*[S3A]*^) system while knocking down *REPTOR* in muscles. Control guts drive *LexAop-nls-sfGFP* (B). DNA is labeled with DAPI (cyan). *REPTOR* knockdown in muscles does not affect tumor development. (**E, F)** TAG (E) and glucose (F) content in thoraces when *REPTOR* expression is reduced in wildtype muscles after 12 days of induction (*dMef2*^*[TS]*^*>REPTOR RNAi*). The reduction of *REPTOR* expression in wildtype muscles does not change TAG or glucose content in thoraces. TAG and glucose were calculated from the same samples. **(G)** Gene expression analysis of the glycolytic rate-limiting enzymes *hex-a* and *hex-c* in *Esg*^*[TS]*^*>yki*^*[S3A]*^ thoraces after 12 days of tumor induction. **(H, I)** Gene expression analysis of *REPTOR* (H) and *pfk* (I) under nutritional restriction. Values were normalized to the mean of control samples (E-G) or fed control samples (H, I). Statistical analysis was done by using one-way ANOVA with Sidak correction test for multiple comparisons (H, I, p<0.05) or using two-tailed *t*-test with Welch’s correction (E-G, p<0.05). Data shows mean with SD.

**Fig. S5:**
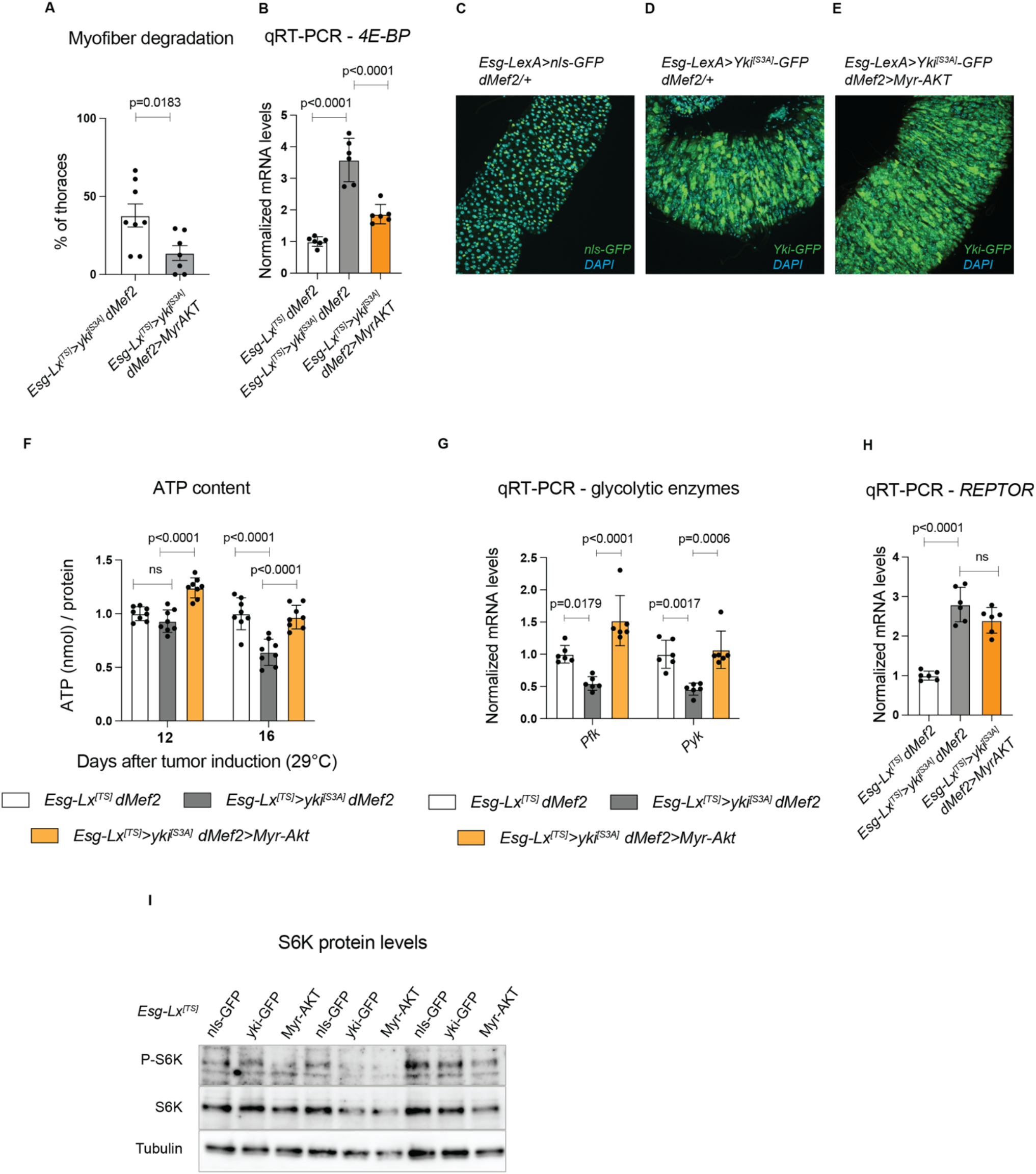
Increasing insulin signaling in *Esg*^*[TS]*^>*yki*^*[S3A]*^ thoraces ameliorates the myofiber degradation but does not modulate REPTOR expression. **(A)** Percentage of thoraces showing myofiber degradation after 19 days of tumor induction. See Table S1. Only *Esg*^*[TS]*^*>yki*^*[S3A]*^ and *EsgLx*^*[TS]*^*>yki*^*[S3A]*^ + *dMef2*^*[TS]*^*>Myr-AKT* are being shown. **(B)** mRNA levels of *4E-BP* in *Esg*^*[TS]*^*>yki*^*[S3A]*^ thoraces with simultaneous muscle-specific *(dMef2*^*[TS]*^*-GAL4)* overexpression of *Myr-AKT* after 12 days of tumor induction. Increased insulin signaling in muscles partially suppresses *4E-BP* upregulation. (**C-E)** Immunostaining of posterior midguts after 12 days of tumor induction with the *EsgLx*^*[TS]*^*>yki*^*[S3A]*^ system while overexpressing *Myr-AKT* in muscles. Control guts drive *LexAop-nls-sfGFP* (B). DNA is labeled with DAPI (cyan). Overexpression of *Myr-AKT* in muscles does not affect tumor development. **(F)** ATP content in thoraces. **(G)** Gene expression analysis of the *pfk* and *pyk* glycolytic rate-limiting enzymes in *Esg*^*[TS]*^*>yki*^*[S3A]*^ thoraces after 12 days of tumor induction. Increasing insulin signaling in muscles of *Esg*^*[TS]*^*>yki*^*[S3A]*^ thoraces upregulates *pfk* and *pyk*. **(H)** mRNA levels of *REPTOR* in *Esg*^*[TS]*^*>yki*^*[S3A]*^ thoraces with simultaneous muscle-specific *(dMef2*^*[TS]*^*-GAL4)* overexpression of *Myr-AKT* after 12 days of tumor induction. *REPTOR* expression is not affected by increased insulin signaling in muscles. **(I)** Protein levels of p-S6K and total S6K from control (*EsgLx*^*[TS]*^*>nls-GFP*), *yki-GFP* (*EsgLx*^*[TS]*^*>yki*^*[S3A]*^) and *Myr-AKT* (*EsgLx*^*[TS]*^*>yki*^*[S3A]*^ + *dMef2*^*[TS]*^*>Myr-AKT*) thoraces. No significant changes in p-S6K were detected when insulin signaling is increased in muscles. Three biological replicates are shown. Values were normalized to the mean of control samples (B, F-H). Statistical analysis was done by one-way ANOVA (B, F-H) with Sidak correction test for multiple comparisons (p<0.05) or using two-tailed *t*-test with Welch’s correction (A, p<0.05). Data shows mean with SD.

**Fig. S6:**
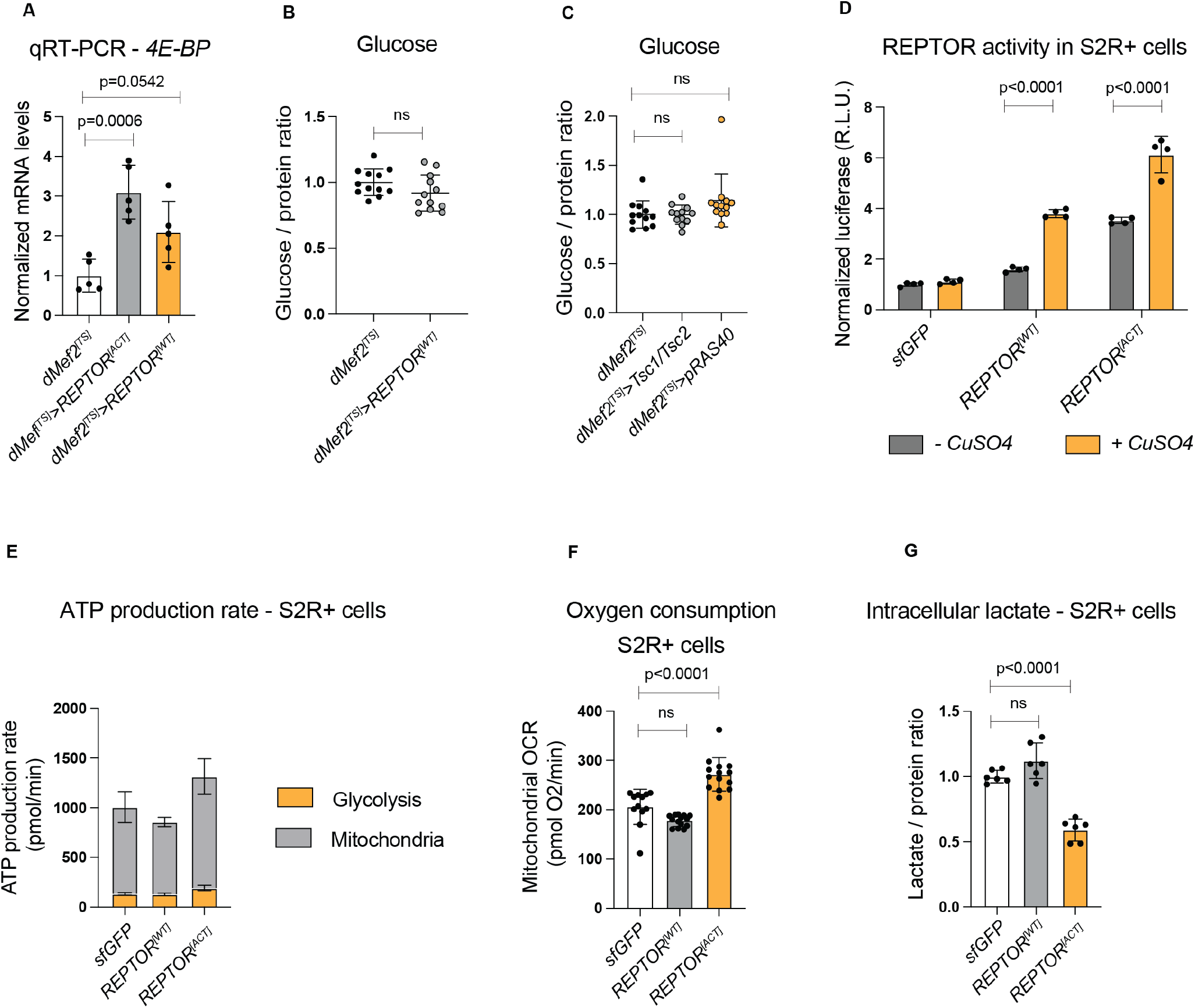
Characterization of the metabolic role of REPTOR in wildtype muscles and in S2R+ cells. **(A)** mRNA levels of *4E-BP* when either an active allele of *REPTOR* (*REPTOR*^*[ACT]*^) or a wildtype allele (*REPTOR*^*[WT]*^) is overexpressed in muscles (*dMef2*^*[TS]*^*-GAL4*) for 4 days. The *REPTOR*^*[WT]*^ mildly upregulates *4E-BP*. (**B, C)** Glucose content in thoraces after 20 days of overexpression of *REPTOR*^*[WT]*^ (B), or of either *Tsc1/Tsc2* or *pRAS40* (C) to reduce mTOR signaling in muscles. No changes in glucose content were detected. **(D)** *Unkempt* luciferase reporter activity in S2R+ cells stably expressing *sfGFP, REPTOR*^*[WT]*^ or *REPTOR*^*[ACT]*^ by induction with CuSO_4_. Luciferase levels are a readout for REPTOR activity. *REPTOR*^*[ACT]*^ S2R+ cells have a leaky expression without induction with CuSO_4_. Luciferase values were normalized to *renilla* luciferase. **(E)** ATP production rate in S2R+ cells *REPTOR*^*[ACT]*^ reduces glycolysis to undetectable levels without changing the contribution of oxidative metabolism (mitochondria). Values for glycolysis and mitochondrial respiration were calculated from twelve to fourteen individual wells for each condition and combined from two different experiments. **(F)** Oxygen consumption in S2R+ cells stably expressing *sfGFP, REPTOR*^*[WT]*^ or *REPTOR*^*[ACT]*^ by induction with CuSO_4_. **(G)** Intracellular amount of lactate in S2R+ cells. Cells expressing *REPTOR*^*[ACT]*^ have significantly less lactate. Values were normalized to the mean of control samples (A-C). The firefly/renilla luciferase ratio was normalized to the average of control samples (*sfGFP* without CuSO_4_) (D). Statistical analysis was done by two-way ANOVA (D) or one-way ANOVA (A, C, F, G) with Sidak correction test for multiple comparisons (p<0.05) or using two-tailed *t*-test with Welch’s correction (B, p<0.05). Data shows mean with SD.

**Fig. S7:**
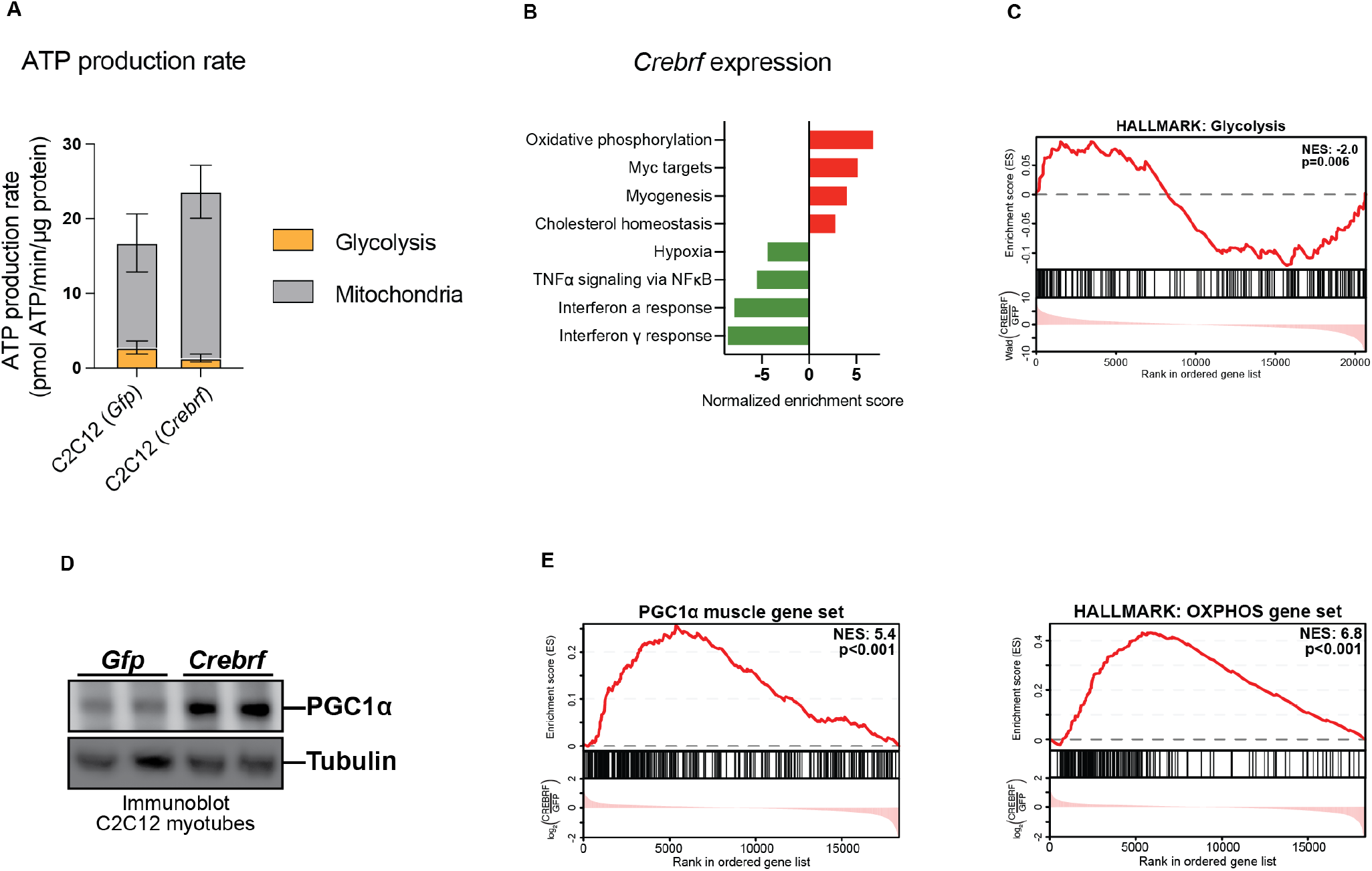
Characterization of the gene expression program driven by CREBRF in mammalian myotubes. **(A)** ATP production rate contributed by glycolysis and mitochondrial respiration in C2C12 myotubes upon adenoviral expression of *Crebrf*. No glucose, pyruvate, or glutamine were present in the media. The ability of CREBRF to repress glycolysis and increase mitochondrial respiration does not depend on exogenous energy substrates. Values for glycolysis and mitochondrial respiration were calculated from ten individual wells for each condition **(B)** Top four HALLMARK gene sets enriched amongst genes up- or down-regulated upon *Crebrf* expression in C2C12 myotubes, as determined by RNA-seq and Gene Set Enrichment Analysis (p<0.001 for each gene set). **(C)** Gene set enrichment plot assessing the HALLMARK glycolysis gene set in C2C12 myotubes expressing *Crebrf*, as measured by RNA-seq. **(D)** Immunoblot of PGC1α protein in C2C12 myotubes expressing *Gfp* or *Crebrf*. TBP immunoblot of input. **(E)** Gene set enrichment plots assessing the expression of the PGC1α muscle gene set (Table S6) or the HALLMARK oxidative phosphorylation gene set. Genes detected in RNA-seq were ranked based on Wald statistic, with the first ranked gene corresponding to the gene most significantly upregulated in *Crebrf* cells. Hash marks represent positions in the ranked list corresponding to members of a given gene set. Normalized enrichment score (NES) indicates whether these members are enriched toward the up-regulated end of this list (positive NES) or the downregulated end of this list (negative NES) as compared to chance expectation. Data shows mean with SD (A).

## Materials and Methods

### *Drosophila* stocks

Bloomington *Drosophila* Stock Center (BDSC): *esg*.*LexA::GAD* (*P{ST*.*lexA::HG}SJH-1*, 66632); *UAS-p60* (*P{UAS-Pi3K21B*.*HA}2*, 25899); *UAS-Myr-Akt* (*P{UAS-myr-Akt*.*ΔPH}3*, 80935); *tub-GAL80*^*[TS]*^ (P{tubP-GAL80[ts]}10, 7108; P{tubP-GAL80ts}7, 7018). National Institute of Genetics Stock Center (NIG): *UAS-ImpL2-RNAi* (15009R-3), *UAS-REPTOR-RNAi* (13624R-3). Vienna *Drosophila* Research Center (VDRC): *Act88f::GFP* (PBac{fTRG10028.sfGFP-FT}, v318362). Laboratory stocks: *w*^*[1118]*^; *tub-GAL80*^*[ts]*^, *dmef2-GAL4; tub-GAL80*^*[ts]*^, *lpp-GAL4; esg-GAL4, UAS-GFP, tub*-*GAL80*^*[ts]*^(*1*); *UAS-yki*^*[S3A]*^(*2*); *UAS-s*.*ImpL2*(*3*); UAS-*Tsc1/Tsc2*(*4*); *UAS-Transtimer*(*5*). The stocks *UAS-REPTOR*^*[ACT]*^, *UAS-REPTOR*^*[WT]*^ and *UAS*.*pRAS40* were a gift from Aurelio Teleman(*6*). *UAS-HA-FoxO* flies were made in this work.

### Crosses

Flies were kept on standard cornmeal fly food supplemented with agar and yeast. Around 20 virgin females were used per cross. Crosses were done under similar conditions of food and kept at 18°C. Bottles were flipped every 4-6 days. For gut *yki*-tumors, males were collected every 24-48 hours and then incubated at 29°C (*GAL4*) or 27°C (*LexAop*) for induction of *yki*^[S3A]^ expression in the gut. For muscle-specific (*dMef2*^*[TS]*^*-GAL4*) or fat body-specific (*lpp*^*[TS]*^*-GAL4*) gene expression, adult males were collected every 48-72 hours, kept at 18°C for an additional 3-4 days and then incubated at 29°C to induce gene expression. Up to 17-18 males were kept per vial to avoid overcrowding and vials at 29°C were flipped every 2-3 days. For a list of genotypes see Table S7.

### Nutritional restriction

Adult males were collected every 48 hours, kept for 5 days at 18°C and then incubated at 29°C for 24 hours to induce gene expression prior to starvation. Flies were then transferred to vials with agarose 0.8% supplemented with 2% sucrose (Fisher Scientific) and maintained at 29°C for 3 days.

### DNA and plasmids

PCR amplification of DNA from plasmids or fly genomic DNA was done using Phusion polymerase (NEB – M0530). Digestion of plasmids with restriction enzymes was done at 37 °C for 2-3 hours. Linearized plasmids and PCR fragments were gel purified using QIAquick columns (QIAGEN – 28115). PCR fragments and plasmid backbones were assembled using Gibson assembly (NEB – E2611). For Gateway cloning, Gateway-compatible expression and entry vectors were recombined using LR Clonase II (Thermo Fisher Scientific – 11791020). Plasmid DNA was extracted from bacterial cultures using a QIAprep Spin Miniprep Kit (QIAGEN – 27104) and Sanger sequenced at the Dana-Farber/Harvard Cancer Center DNA Resource Core or Genewiz. The *unkempt-*luciferase plasmid(*6*) was a gift from Aurelio Teleman and the *act5c-renilla(7)* was a gift from Nirmalya Chatterjee.

### Generation of the *LexAop-nls-sfGFP* and *LexAop-yki*^*[S3A]*^ fly stocks

The *nls-sfGFP* construct was amplified from a T2A-sfGFP plasmid kindly provided by Justin Bosch (unpublished) with the following oligos: forward primer “CTTCAGGCGGCCGCGGCCAACATGCCCAA-GAAGAAGCGCAAGGTGGTGTCCAAGGGCGAGGAGC” and reverse primer “CTTCACAAAGATCCTCTAGCTACTTGTACAGCTCATCCATG”. The *yki*^*[S3A]*^ sequence was amplified from genomic DNA of *UAS-yki*^*[S3A]*^ flies using as forward primer “GTACGAATTCCAACATGTTAACGACGATGTCAGCC” and the reverse primer “ATATGCGGCCGCTCACGTAGAATCGAGAC” that includes an homologous sequence to the V5-tag present in the *yki*^*[S3A]*^ transgene(*2*). The *yki*^*[S3A]*^ PCR fragment was cloned in frame with *sfGFP* (Justin Bosch - unpublished). The *yki*^*[S3A]*^*-sfGFP* fragment was amplified using the following oligos: forward primer “CCTTTACTTCAGGCGGCCGCGGCCACCATGTT-AACGACGATGTCA” and reverse primer “CTTCACAAAGATCCTCTAGCTA-CTTGTACAGCTCATCCATG”. The *nls-sfGFP* and *yki*^*[S3A]*^*-sfGFP* PCR products were then cloned by Gibson assembly in the backbone of the pJFRC19-*13XLexAop2-IVS-myr::GFP* plasmid (Addgene – 26224) previously digested with XbaI and XhoI to remove the *myr::GFP* sequence. Final plasmids were injected in fly embryos of the stock *yw; nos-Cas9 attP40* for targeted insertion in the second chromosome.

### Generation of *UAS-HA-FoxO* line

The complete cDNA of *dFOXO* was obtained from the BDGP collection, clone LD19191 (accession no. AF426831), and subcloned into a gateway entry vector using the following oligo sequences: forward primer “CACCATGATGGACGGCTACGCGCAGGAATG” and reverse primer “CTAGTGCACCCAGGATGGTGGCGAGGTCAC”. The Akt phosphorylation sites in dFOXO were mutated to T44E, S190E, and S259E using the oligos sequences: forward/reverse primers “CGGGCCAGATCCAACGAGTGGCCATGTCCGCG”/ “CGCGGACATGGCCA-CTCGTTGGATCTGGCCCG” for T44E; “CGCCGCCGTGCCGCTGAGATGGAG-ACGTCCCGG”/ “CCGGGACGTCTCCATCTCAGCGGCACGGCGGCG” for S190E; and “CGGCAACGCGCCTCAGAGAATGCCAGTTCCTGC”/ “GCAGGAACTGGCATTCTCTGAGGCGCGTTGCCG” for the S259E position, according to the manufacturer’s procedure for the multisite mutagenesis kit (Invitrogen). The resulting construct was cloned into the pHWUASt destination vector from the Drosophila Gateway Vector collection to generate a N-terminal HA-tagged *FoxO*. Transgenic flies were obtained by injecting the construct into *w*^*-*^ recipients (Bestgene). *UAS-HA-FoxO* (2^nd^ chromosome) flies were then crossed to a *dMef2*^*[TS]*^*-GAL4* to confirm the constitutive nuclear localization of FoxO in muscles by immunostaining with an anti-HA antibody.

### Protein, TAG and glucose colorimetric measurements

Protocol was adapted from(*8*). Five or six biological replicates were collected from each cross at the designated time point. For each biological sample, eight thoraces were dissected in PBS by clipping the wings and the head, separating the abdomen from the thorax and making sure the entire gut was removed. Dissected thoraces were immediately frozen on dry ice and stored at -80°C until collections were finished. Samples were placed in an Abgene™ 96 Well 1.2mL Polypropylene Deepwell Storage Plates (Thermo Fisher Scientific – AB0564) with 200ul of ice-chilled PBS with 0.05% Triton-X-100 (Sigma) and Zirconium Oxide Beads, 1.0mm, 1Lb. (Next Advance Lab Products – ZROB10). Homogenization was done using a TissueLyser II homogenizer (QIAGEN), 2-3 times per 30 seconds with an oscillation frequency of 30Hz/second. Plates were then centrifuged for 1 min at 4000 rpm to spin down debris and the homogenate was immediately used to quantify glucose and protein. For TAG, an additional step of heat treatment for 10’ at 75°C was performed. Quantifications were done using a SpectraMax Paradigm Multi-mode microplate reader. For protein measurements, 5 ul of homogenate was diluted in 20ul of PBS and mixed with 200ul of BCA Protein Assay Kit (Pierce™ BCA Protein Assay Kit), and samples were incubated for 30 minutes at 37°C with gentle shaking in 96-well Microplates (Greiner Bio-One). Glucose was measured by mixing 10 ul of homogenate in 100ul of Infinity Glucose Hexokinase Reagent (Thermo Fisher Scientific™ - TR15421), and samples were incubated for 30 minutes at 37°C in 96-well Microplates UV-Star (Greiner Bio-One – 655801) with gentle shaking. For TAG measurements, 20 ul of homogenate were used in 150 ul of Triglycerides Reagent (Thermo Fisher Scientific™ - TR22421) and incubated for 10 minutes at 37°C in 96-well Microplates (Greiner Bio-One) with gentle shaking. Values were determined from standard curves of 5 serial dilutions (1:1) of BSA (Pierce™ BCA Protein Assay Kit), Glycerol standard solution (Sigma) or Glucose standard solution (Sigma). All experiments were independently performed at least twice.

### RNA extraction and RT-qPCR

*<u>Drosophila</u>* <u>thoraces and guts</u>: thoraces were dissected as described above. Five to ten thoraces or six to ten guts were collected per sample. Samples were homogenized in TRIzol reagent (Ambion) and mixed vigorously with Chloroform (Fisher) for phase separation. The collected supernatant was processed on Direct-zol™ RNA MicroPrep columns (Zymo Research) according to the manufacturer instructions. Eluted RNA was treated with TURBO DNA-free™ Kit (Ambion) and cleaned up again with the Direct-zol™ RNA MicroPrep columns. Same amount of total RNA was used for reverse transcription by using iScript™ cDNA Synthesis Kit (Bio-Rad). RT-qPCR was done in a CFX96 Real-Time System (Bio-Rad) using iQ™ SYBR® Green Supermix (Bio-Rad). Relative mRNA levels were calculated using the ΔΔCt method. For thoraces, values were normalized to *a-tubulin* unless stated to *rp49*; for guts, mRNA levels were normalized to *rp49*. <u>Mouse C2C12 myotubes</u>: Total RNA was extracted using TRIzol and purified using RNeasy Mini spin columns (QIAGEN). Reverse transcription was performed using a High-Capacity cDNA Reverse Transcription kit (Applied Biosystems). The resultant cDNA was assessed by RT-qPCR using SYBR® Green Supermix 2x qPCR master mix (Promega) in a ABI PRISM 7900HT Real-Time PCR system (Applied Biosystems). Relative mRNA levels were normalized to *Tbp* mRNA and calculated using the ΔΔCt method(*9*). Primer sequences are listed in Table S8.

### RNA-Seq

<u>Library preparation for transcriptome analysis</u>: Total RNA was quantified using an Agilent 2200 or 4200 TapeStation instrument, with a corresponding Agilent TapeStation RNA assay. The resulting RIN (RNA Integrity Number) scores and concentrations were considered for qualifying samples to proceed, with RIN spanning between 8.8 and 9.6. For the *Esg*^*[TS]*^*>yki*^*[S3A]*^ time-course experiment, the rRNA was depleted using Illumina RiboZero HMR workflow. For the REPTOR and CREBRF overexpression experiments, the mRNA was pulled-down using oligo-dt beads as part of the KAPA mRNA HyperPrep workflow. Library samples were normalized in equimolar ratio and divided into four pools for technical replication. Each pool was denatured and loaded onto an Illumina NextSeq 500 instrument, with a High-Output 75-cycle kit to obtain Single-Read 75bp reads. <u>Transcriptome analysis –</u> *<u>Drosophila</u>*: FastQC was used for sequencing data quality control and MultiQC was used to aggregate the results. Sequencing reads were aligned to Drosophila reference genome obtained from FlyBase (FB dmel r6.30) using STAR. DSEQ2 is used to identify the differentially expressed genes using biological replicates or technical replicates from time series experiment and final hits were selected based on both adjusted P value as well as fold change cut-off. 510 genes were selected from the RNA-seq dataset time-course induction of *yki* tumors using 2.5-fold change cut-off, while 700 gene hits were selected from the RNA-seq dataset of REPTOR overexpression in muscle using 2.5-fold change cut-off with adjust p value <0.05. The information of differentially expressed genes of each data set are presented in Table S2 and S4 along with gene ontology annotation and respective human orthologs identified by DIOPT. TFs were selected from differentially expressed genes of time-course RNA-seq dataset and were illustrated by heatmap using TM4 software suite (http://mev.tm4.org/). TF annotation was obtained from GLAD (PMID: 26157507). Enrichment analysis was performed on differentially expressed genes from both datasets using an in-house program based on hyper-geometric distribution. The gene sets were assembled based on the gene ontology (GO) annotation from FlyBase and SLIM terms specifically for Drosophila genes were selected. KEGG pathway annotation for Drosophila were selected and supplemented with the annotation for human genes that were mapped to Drosophila using DIOPT with the high/moderate rank filter. Enrichment results for each dataset set are presented in Supplementary Tables 3 and 5. <u>Transcriptome analysis –</u>*<u>Mouse C2C12 myotubes</u>*: Sequenced reads were trimmed using fastp 0.20.1(*10*) and then aligned to the mm10 reference genome assembly using RNA STAR 2.7.8(*11*). Differential gene expression testing was performed by analyzing gene counts from RNA STAR using DESeq2 2.11.40.6(*12*). Genes with mean expression below 0.5 were filtered, and the remaining genes were ranked by Wald statistic to generate a pre-ranked gene list for gene set enrichment analysis using GSEA v4.0.1(*13, 14*). ‘Classic’ mode was used for enrichment statistic calculation. For unbiased discovery of enriched gene sets, the HALLMARK gene sets were used as queries(*15*). In a separate query, muscle gene expression data were examined using a custom gene set: ‘PGC1α muscle gene set.’ (Table S6). This gene set consists of 331 genes upregulated in the gastrocnemius muscle of MCK-PGC1α transgenic mice with a p value <1e-5(*13*). P values reported in figures correspond to FDR q values to account for multiple hypothesis testing.

### Immunostaining of thoraces and guts

Thoraces were fixed for 30 minutes on a relaxing buffer composed by PBS with 0.3% Triton-X-100 (PBT 0.3%), MgCl2 [5mM], EGTA [5mM], ATP [5mM] and 4% formaldehyde (Polysciences). After fixation, thoraces were bisected with a Double Edge Razor Blade (Personna – 74-0002), washed three times in PBT 0.3% and then incubated in PBT 0.3% with 5% BSA for 1 hour at room temperature. For gut immunostaining, dissected guts were fixed on PBT 0.1% with 4% formaldehyde for 20 minutes, washed three times in PBT 0.1% and then blocked in PBT 0.1% with 5% BSA for 1 hour. Incubation with primary antibodies was done overnight at 4°C in blocking solution. Antibodies used were mouse anti-CoxIV (1:500, Abcam – 33985), chicken anti-GFP (1:2000, Aves Labs – GFP-1020) and anti-bPS1 (1:50, DSHB – CF.6G11). Corresponding secondary antibodies used were Alexa Fluor™ 488 or Alexa Fluor™ 647 (1:500, Jackson Immunology). For staining of myofibrils, Alexa Fluor™ 555 Phalloidin (1:200, Thermo Fisher Scientific) was incubated with secondary antibodies for 2 hours at room temperature. Hemithoraces were mounted on u-Dish 35mm high Glass Bottom (Ibidi) in VECTASHIELD® Antifade Mounting Medium with DAPI (Vector Laboratories)

### Thorax cross-sections

To make agarose blocks, Intermediate Tissue-Tek Cryomolds (Sakura – 4566) were loaded with agarose 1% and cooled down. Three intact thoraces previously fixed in PBT 0.3% with 4% formaldehyde for 20 minutes were placed in the agarose before solidifying. Slices of the agarose blocks with around 300um thickness were generated on a Vibratome (Leica VT 1000M). Samples were carefully collected and immediately incubated in a blocking solution of PBT 0.3% with 5% BSA for 1 hour at room temperature and then followed by immunostaining.

### Confocal imaging and analysis

<u>Confocal imaging</u> was performed using either: i) fully motorized Nikon Ti or a fully motorized Ti2 inverted microscope equipped with a Yokogawa CSU-W1 Spinning disk scanhead with a single 50 um pinhole disk, a PI 250 um or Mad City Labs 500 um Z piezo stage inserts, a TOPTICA iChrome MLE laser launch with directly modulated 405nm 100mW solid state, 488 nm solid-state, 561 nm DPSS and 640 nm solid state laser lines, and a SOLA SE Lumencor light engine for widefield illumination. Image acquisition was done using either a Plan Apo λ 60X 1.4 NA immersion oil DIC used with Cargille Type 37 immersion oil (Figure 1f; Figure 3b), a Plan Fluor 40x/1.3 NA immersion oil used with Cargille Type 37 immersion oil (Extended Data Fig. 4b-d; Extended Data Fig. 5b-d), a 20x/0.75 N.A. Air DIC objective (Figure 1d, e; Figure 3b) or a Plan Apo 10x/0.45 Air DIC objective (Figure 1A). Images were captured using an Andor Zyla 4.2 Plus sCMOS monochrome camera using the 16-bit dual gain digitizer mode and Nikon Elements Acquisition Software AR 5.02. Signal from the different channels was acquired sequentially using a Semrock Di01-T405/488/568/647 multi-band pass dichroic mirror and band pass emission filters for green (Chroma ET525/36m), red (Chroma ET 605/52m), far read (Chroma ET 705/72m) channels and DAPI (Chroma ET455/50m). Z stack acquisition was optimized for proper axial sampling, 0.3 um step size for 60x, acquiring all channels in the same focal plane to minimize axial shifts. For 40x objective lens (0.3 um step size), 20x objective lens (0.9 um step size) and 10x objective lens (5.6 um step size) each channel was acquired sequentially. ii) a Zeiss LSM780 single point scanning confocal unit outfitted with galvanometer mirrors attached to a fully motorized Axio-Observer Z1 equipped with a Zeiss motorized stage. Samples were excited sequentially by illuminating with a 405 nm diode 30 mW AOTF modulated line set to 0.2% transmittance, a 488 nm 20 mW Argon multi-line AOTF modulated line set to 0.4% transmittance, and a 561 nm 25 mW DPSS AOTF modulated line set to 0.2% transmittance. Images were acquired using ZEN Black SP2 acquisition software, scanned bi-directionally with a pixel dwell time of 1.27 µsec/pixel, line average of 2, 8-bit digitizer and pinhole diameters corresponding to 1 A.U. for each wavelength. Each Z stacks was acquired using a Zeiss Plan Apochromat 63x/1.4 NA DIC oil immersion objective lens with Zeiss Immersol 518 F immersion oil, pixel size of 0.13×0.13 µm2 (1024×1024 frame size and no zoom) and using 0.3um step size (Extended Data Fig. 1 a, b), or a Zeiss LD LCI Plan Apochromat 25x/0.8 multi-immersion lens with correction collar set up to oil, 0.4 um/pixel (512×512 frame size and no zoom) and using 1 um step size (Extended Data Fig. 1 c, d). <u>Image analysis:</u> was done in Fiji/ImageJ(*16*). For images of hemithoraces acquired with the 60x or 63x objective, maximum intensity projections were done using 10-15 stacks of each acquisition starting from the surface of the sample and a representative image was shown. For the 10x and 20x scans of hemithoraces and 40x scans of posterior midguts, the maximum intensity projections included all stacks. For cross-sections, maximum intensity projections were done using 10-15 slices starting from the surface of the sample.

### Myofiber degradation analysis

For each experiment, up to 15 flies per vial were incubated at 29°C and then thoraces were dissected at a defined time point, fixed for 40 minutes in relaxing buffer, bisected with a razor blade, and stained for 2 hours with Alexa Fluor™ 555 Phalloidin (1:200, Thermo Fisher Scientific – A34055) at room temperature. Hemithoraces were mounted on u-Dish 35mm high Glass Bottom (Ibidi) in VECTASHIELD® Antifade Mounting Medium (Vector Laboratories). Z stack acquisition of samples on the Yokogawa CSU-W1 Spinning disk was done with the 20x objective and each hemi-thorax was analyzed for the presence of myofiber degradation in any of the dorso-longitudinal muscles.

### ATP quantification

ATP quantification protocol was adapted from(*17*). Four biological replicates, each with 4 thoraces homogenized in 100ul of 5M guanidine-HCl (Sigma) supplemented with 100 mM Tris and 5 mM EDTA, pH 7.8, immediately boiled for 3 minutes at 95°C, centrifuged, and then the supernatant collected. Extracts were used for quantification of luciferase as proxy for ATP content using the ENLITEN® ATP Assay System Bioluminescence Detection Kit (Promega) according to the manufacturer instructions. Samples were diluted in water to fit the range of the standard curve (5 serial dilutions, 1:10), and ATP levels were calculated from the standard curve by measuring the luminescence in a SpectraMax Paradigm Multi-mode microplate reader using Corning® 96-well Solid White Flat Bottom Polystyrene microplates (Costar). Values were normalized to protein content (Pierce™ BCA Protein Assay Kit). All experiments were independently performed three times.

### Data analysis, statistics and reproducibility

GraphPad Prism was used for statistical analysis. For analysis of two groups, unpaired two-tailed t-test with Welch’s correction was used to calculate the p-value (p<0.05). For analysis of three groups, two-way ANOVA or one-way ANOVA were used with Sidak multiple comparisons test to calculate the p-value (p<0.05). All data values were normalized to the mean of control samples. All experiments were independently repeated at least twice. were generated by combining the values of two independent experiments. No randomization or blinding was done. Quantifications of the myofiber degradation phenotypes are in the Table S1. Complete information on genotypes is detailed in Table S7. List of qPCR primers and their source is listed in Table S8.

### Western Blot

*<u>Drosophila thoraces</u>*: Around 10 thoraces were homogenized in RIPA Lysis and Extraction Buffer (Thermo Fisher Scientific) with Halt™ Protease and Phosphatase Inhibitor Cocktail (100X) (Thermo Fisher Scientific), incubated for 10 minutes on ice and then centrifuged for 15 minutes at 15000 rpm at 4°C. 5 ul of each supernatant sample was used to quantify the total amount of protein with a BCA Protein Assay Kit whereas the rest was diluted in 4x Laemmli sample buffer (Bio-Rad) with 2-Mercaptoethanol (Sigma), and then boiled for 10 minutes at 90°C. Samples were further resolved on Mini-PROTEAN® TGX™ Precast Protein Gels (Bio-Rad) and transferred to Immobilon-P PVDF Membrane (Millipore IPVH00010). Membranes were blocked on a 5% dry nonfat milk solution or 5% BSA with Tris-buffered saline (TBS) with 0.1% TWEEN® 20 (Sigma) at room temperature for 1 hour. Further incubation with primary antibodies was done o/n at 4°C with gentle shaking. After washing, membranes were incubated with HRP-conjugated secondary antibodies at room temperature for 1 hour and developed using a SuperSignal™ West Pico PLUS Chemiluminescent Substrate (Thermo Fisher Scientific – 34580). Primary antibodies used were rabbit anti-phospho AKT (1:1000, Cell Signaling – 4054), rabbit anti-Akt (1:1000, Cell Signaling – 9272), rabbit anti-pS6K (1:1000, Cell Signaling – 9209), total anti-S6K(*18*), mouse anti-tubulin (1:5000, Sigma – T5168). <u>Mouse C2C12 myoblasts</u>: Cell culture samples were prepared in ice-cold RIPA buffer (50 mM Tris pH 7.4, 150 mM NaCl, 1% NP40, 0.5% sodium deoxycholate, 0.1% SDS) supplemented with 1x Complete EDTA-free protease inhibitor (Roche) and 1x Phos-stop phosphatase inhibitor (Roche). Lysates were sonicated 7.5 min in a Diagenode water bath sonicator (high intensity; 30 s on, 30 s off cycles), centrifuged at 16,000 g for 15 min at 4 °C, and the supernatants were used for subsequent analyses. Protein concentration was determined using the BCA assay. Protein lysates were denatured in Laemmli buffer, resolved by 4-12% NuPAGE Bis-Tris SDS-PAGE (Invitrogen) and transferred to polyvinylidene difluoride (PVDF, 0.45 μm pore size) membrane. Primary antibodies were diluted in TBS containing 0.05% Tween-20, 5% BSA, and 0.05% NaN3. Primary antibodies were: rabbit anti-TBP (1:1000, Cell Signaling – 44059) and mouse anti-PGC1α (1:1000, EMD Millipore – ST1202). Membranes were incubated overnight with primary antibodies at 4°C. For secondary antibody incubation, anti-rabbit or anti-mouse HRP (Promega) was diluted in TBS containing 0.05% Tween-20 and 5% dry nonfat milk. HRP signal was visualized using Crescendo Western HRP substrate (EMD Millipore) and an Amersham Imager 680. Detection of PGC1α required immunoprecipitation prior to immunoblotting. After preparation of RIPA lysates as described above, 0.5 mg of protein was immunoprecipitated using 5 μg anti-PGC1α antibody (Santa Cruz – sc518025) during overnight rotation at 4°C. The next day, 20 μl of protein G agarose bead slurry (Thermo Fisher) was washed and added to each IP, followed by rotation at 4°C for 2 hr. Beads were washed 3x in RIPA buffer and immunoprecipitated protein was eluted in Laemmli buffer for analysis by immunoblot. Immunoblot was performed as above, except that the anti-PGC1α primary antibody was diluted in TBS with 0.05% Tween-20 and 5% nonfat dry milk during overnight incubation.

### Cell culture

<u>Drosophila S2R+ cells</u>: Drosophila S2R+ cells were cultured at 25°C in Schneider’s *Drosophila* medium (Thermo Fisher Scientific), supplemented with 10% fetal bovine serum (Sigma) and 50 U/mL of penicillin-streptomycin (Thermo Fisher Scientific). For transfections of S2R+ cells with plasmids, Effectene Transfection Reagent (QIAGEN) was used following the manufacturer’s instructions. To generate S2R+ cells stably expressing *REPTOR*^*[ACT]*^ or *REPTOR*^*[WT]*^, the wildtype sequence of *REPTOR* was amplified from genomic DNA of w^*[1118]*^ flies using the following oligos: forward primer “CCGCGGCCGCCCCCTTCACCATGACAGAGAATCAGCTGTA” and reverse primer “GGGTCGGCGCGCCCACCCTTCATATAAAGCCCAGGCTCTT” and then cloned in a gateway-compatible entry vector by Gibson assembly reaction. The REPTOR^*[ACT]*^ allele was generated by doing an additional PCR amplification step of site-directed mutagenesis on the *REPTOR*^*[WT]*^ sequence to mutate the two serine aminoacids (S527 and S530) to alanine using the following oligos: forward primer GAGCCACGACAGCATTTACTCGC-CACCGGCGCGCTGGCCGAGGCCGAGAGCTTCTCTTC” and reverse primer “GAAGAGAAGCTCTCGGCCTCGGCCAGCGCGCCGGTGGCGAGTAAATGCTGTCGTG GCTC”. The PCR product, which included the full sequence of the entry vector, was treated with DpnI for 1 hour at 37°C and then transformed into Chemically competent TOP10 *Escherichia coli* for plasmid extraction with. The *sfGFP* sequence was amplified from the *LexAop-sfGFP* plasmid and cloned in a gateway-compatible entry vector using Gibson assembly using the following oligos: forward primer “CCGCGGCCGCCCCCTTCACCATGGTGTCCAAGGGCGAG-GAGCTGTTC” and reverse primer “GGGTCGGCGCGCCCACCCTTCTA-CTTGTACAGCTCATCCATG”. Each gene was then cloned in the *pMK33-GW* plasmid(*19*) using gateway cloning. S2R+ cells were transfected with the plasmids pMK-*REPTOR*^*[ACT]*^, pMK-*REPTOR*^*[WT]*^ or pMK-*sfGFP* in normal media. After 72 hours, cells were incubated and expanded in media supplemented with 200 µg/mL hygromycin B (Calbiochem) for 4 weeks. To induce gene expression, CuSO_4_ was added to each well at a final concentration of 100uM. Activity of the *REPTOR*^*[ACT]*^ and the *REPTOR*^*[WT]*^ alleles was measured by co-transfecting by each cell line with the *unk-*luciferase reporter plasmid(*6*) and the *pAct-renilla* plasmid(*7*). After 72 hours, the firefly luciferase signal of the *unk-*reporter and the *renilla* luciferase signal were quantified using the Dual-Glo Luciferase Assay system (Promega) following the manufacturer instructions. Firefly luciferase values were normalized to *renilla*. <u>Mouse C2C12 cells</u>: C2C12 cells (ATCC) and HEK293A cells (Thermo Fisher Scientific) were cultured in DMEM with 4.5 g/L glucose and 584 mg/L L-glutamine (Corning), supplemented with 10% sterile filtered fetal bovine serum (GeminiBio) and 1x penicillin/streptomycin (Gibco). Cells were maintained in 37°C and 5% CO2. Adenoviral vectors encoding *Gfp* or M. musculus *Crebrf* (Uniprot Q8CDG5-1) were constructed using the pAd/CMV/V5-DEST vector (Thermo Fisher). No epitope tag was added to the CREBRF polypeptide. Adenovirus was generated in HEK293A cells and titered using the Adeno-X Rapid Titer Kit (Takara Bio). For differentiation, C2C12 cells were grown to confluence in 6-well cell culture plates and washed with PBS, followed by addition of DMEM with 4.5g/L glucose and 584 mg/L L-glutamine (Corning), supplemented with 2% horse serum (GemCell) and 1x penicillin/streptomycin (Gibco). Medium was changed every two days thereafter. For viral transduction experiments, adenovirus was added to the medium for 3 hours on day 4 of differentiation at 250 MOI in the presence of 5 μg/ml polybrene (Santa Cruz Biotechnology). Experiments were performed 3 days later, at day 7 of differentiation. To assess the effects of nutrient withdrawal, 6 well plates containing C2C12 myotubes differentiated to day 7 as above, were washed once with 3 ml PBS and then incubated in Hank’s balanced salt solution without calcium or magnesium (Corning) for 4 hr. To assess insulin signaling, C2C12 myotubes were differentiated to day 6 as above, washed with 3 ml PBS, and then incubated overnight (16 hr) in DMEM without serum. Bovine pancreatic insulin (Sigma) was added to 100 nM for 10 min and then cells were harvested for immunoblot analysis.

### Mice

Mice were housed at 23°C under a 12 hour light/dark cycle. All experiments used 8-12 week old male C57BL/6 mice (Jackson Labs). Animal experiments were performed according to procedures approved by the Institutional Animal Care and Use Committee (IACUC) of the Beth Israel Deaconess Medical Center.

### Cellular bioenergetic measurements in S2R+ cells and C2C12 myotubes

S2R+ cells were incubated with CuSO4 at 100uM to induce gene expression. After 48 hours of induction, cells were seeded at a density of hundred and fifty thousand cells per well in 24-well Seahorse XF V7-PS cell culture microplates, previously treated with ConcanavelinA (Sigma) at a concentration of 0.1mg/ml overnight at 4°C, and an ATP rate assay was performed as described below. Similarly, C2C12 cells were seeded in 24-well Seahorse XF V7-PS cell culture microplates and differentiated upon confluence, then transduced using adenovirus as described above. Three days after transduction, an ATP rate assay was performed. The ATP rate assay was carried out similarly for Drosophila and mammalian cells. The cells were washed twice with 0.5 ml Seahorse XF DMEM medium pH 7.4 (Agilent) and then incubated in a CO2-free incubator at 37 °C (or 24°C for S2 cells) for 45 minutes. Unless otherwise stated, the medium was supplemented with 10 mM glucose, 1 mM pyruvate, and 2 mM glutamine. At this point, the medium was removed and replaced with 0.5 ml Seahorse XF DMEM. Oxygen consumption and extracellular acidification were assessed in a standard ATP rate assay protocol in a XFe24 Seahorse Analyzer at either 37°C or 24°C as follows: basal measurement (3 cycles), inject port A (3 cycles), inject port B (3 cycles). Injection port A contained oligomycin (1.5 μM final; Cell Signaling Technology) and injection port B contained rotenone/antimycin A (1 μM/1 μM final; Sigma). Each cycle consisted of 2 min 30 sec mixing, 2 min waiting, and 3 min measuring. The contribution of respiration and glycolysis to ATP production were calculated using the Seahorse Analytics web application (Agilent). Measurements were normalized to protein content as determined by BCA assay (Thermo Fisher Scientific).

### Glucose uptake assay

C2C12 myotubes were differentiated in 12-well plates and transduced as described above. At day 6 of differentiation, the cells were washed with PBS and then incubated overnight (16 hr) in DMEM lacking serum. On day 7, 1 μM insulin or vehicle control was added to each well and cells were incubated for an additional 1 hr at 37°C, at which point the medium was removed and replaced with 0.25 ml PBS containing 0.1 mM 2-deoxyglucose. The cells were incubated an additional 30 min at 24°C and then lysed in stop buffer, allowing measurement of 2-deoxyglucose-6-phosphate using the Glucose Uptake-Glo assay (Promega) and a FLUOstar Omega luminescence plate reader.

### Lactate measurement

S2R+ cells were incubated with CuSO4 at 100uM to induce gene expression for 48 hours. Cells then were washed twice with ice-cold PBS and lysed by a mixture of 0.25 ml PBS and 0.125 ml 0.6 N HCl. Plates were rocked at 22°C for 5 min, followed by addition of 0.125 ml 1M Tris base. Intracellular lactate was subsequently measured using the Lactate-Glo assay (Promega) and a FLUOstar Omega luminescence plate reader. For C2C12 myotubes, DMEM medium with 2% horse serum was conditioned for 24 hr by C2C12 myotubes transduced as above with adenovirus encoding *Gfp* or *Crebrf*. The medium was subsequently assessed for lactate content using a Lactate Colorimetric Assay Kit (Biovision).

